# Bioenergetics of human spermatozoa in patients with testicular germ cell tumour

**DOI:** 10.1101/2024.05.24.595824

**Authors:** Ondrej Simonik, Barbora Bryndova, Vishma Pratap Sur, Lukas Ded, Zuzana Cockova, Ales Benda, Maryam Qasemi, Petr Pecina, Alena Pecinova, Daniela Spevakova, Tomas Hradec, Pavel Skrobanek, Zuzana Ezrova, Zuzana Kratka, Radomir Kren, Michal Jeseta, Ludmila Boublikova, Libor Zamecnik, Tomas Büchler, Jiri Neuzil, Pavla Postlerova, Katerina Komrskova

**Affiliations:** Laboratory of Reproductive Biology, Institute of Biotechnology of the Czech Academy of Sciences, Vestec, Czech Republic; Department of Biochemistry, Faculty of Science, Charles University, Prague, Czech Republic; Imaging Methods Core Facility at BIOCEV, Faculty of Science, Charles University, Vestec, Czech Republic; Laboratory of Bioenergetics, Institute of Physiology, Czech Academy of Sciences, Prague, Czech Republic; Department of Urology, General University Hospital and First Faculty of Medicine, Charles University, Prague, Czech Republic; Department of Oncology, First Faculty of Medicine, Charles University and Thomayer University Hospital, Prague, Czech Republic; Laboratory of Molecular Therapy, Institute of Biotechnology of the Czech Academy of Sciences, BIOCEV, Vestec, Czech Republic; Laboratory of Immunology, IVF Clinic GENNET, Prague Czech Republic; Laboratory of embryology, IVF Clinic GENNET, Prague, Czech Republic; Department of Obstetrics and Gynaecology, Faculty of Medicine, Masaryk University and University Hospital Brno, Brno, Czech Republic; Department of Oncology, Second Faculty of Medicine and Motol University Hospital, Prague, Czech Republic; School of Pharmacy and Medical Science, Griffith University, Southport, QLD, Australia; Department of Zoology, Faculty of Science, Charles University, Prague, Czech Republic

**Keywords:** TGCT, spermatozoa, sperm biochemistry, sperm function, energetic metabolism, oxidative phosphorylation, Seahorse, 2P-FLIM, infertility, cancer

## Abstract

In testicular germ cell tumour (TGCT) patients, sperm cryopreservation prior to anti-cancer treatment represents the main fertility preservation approach. However, it is associated with low sperm recovery rate after thawing. Since sperm is a high-energy demanding cell, which is supplied by glycolysis and oxidative phosphorylation (OXPHOS), mitochondrial dysfunctionality can directly result in sperm anomalies. In this study, we investigated the bioenergetic pattern of cryopreserved sperm of TGCT patients in comparison with normozoospermic samples using two state-of-the-art methods; the Extracellular Flux Analyzer (XF Analyzer) and Two-Photon Fluorescence Lifetime imaging (2P-FLIM), in order to assess the contributions of OXPHOS and glycolysis to energy provision. A novel protocol for combined measurement of OXPHOS (Oxygen Consumption Rate – OCR) and glycolysis (Extracellular Acidification Rate – ECAR) using the XF Analyzer was developed together with a unique customized AI-based approach for semiautomated processing of 2P-FLIM images. Our study delivers optimized Low-HEPES modified Human Tubal Fluid media (mHTF) for sperm handling during pre-analytical and analytical phases to maintain sperm physiological parameters and optimal OCR, equivalent of OXPHOS. The negative effect of cryopreservation was signified by deterioration of both bioenergetic pathways represented by modified OCR and ECAR curves and the derived parameters. This was true for normozoospermic as well as TGCT samples, which showed even a stronger damage within the respiratory chain compared to the level of glycolytic activity impairment. The impact of cryopreservation and pathology are supported by 2P-FLIM analysis showing a significant decrease in bound NADH in contrast to unbound NAD(P)H which reflects decreased metabolic activity in samples from TGCT patients. Our study provides novel insight into the impact of TGCT on sperm bioenergetics and delivers a verified protocol to be used for assessment of human sperm metabolic activity, which can be a valuable tool for further research and clinical andrology.

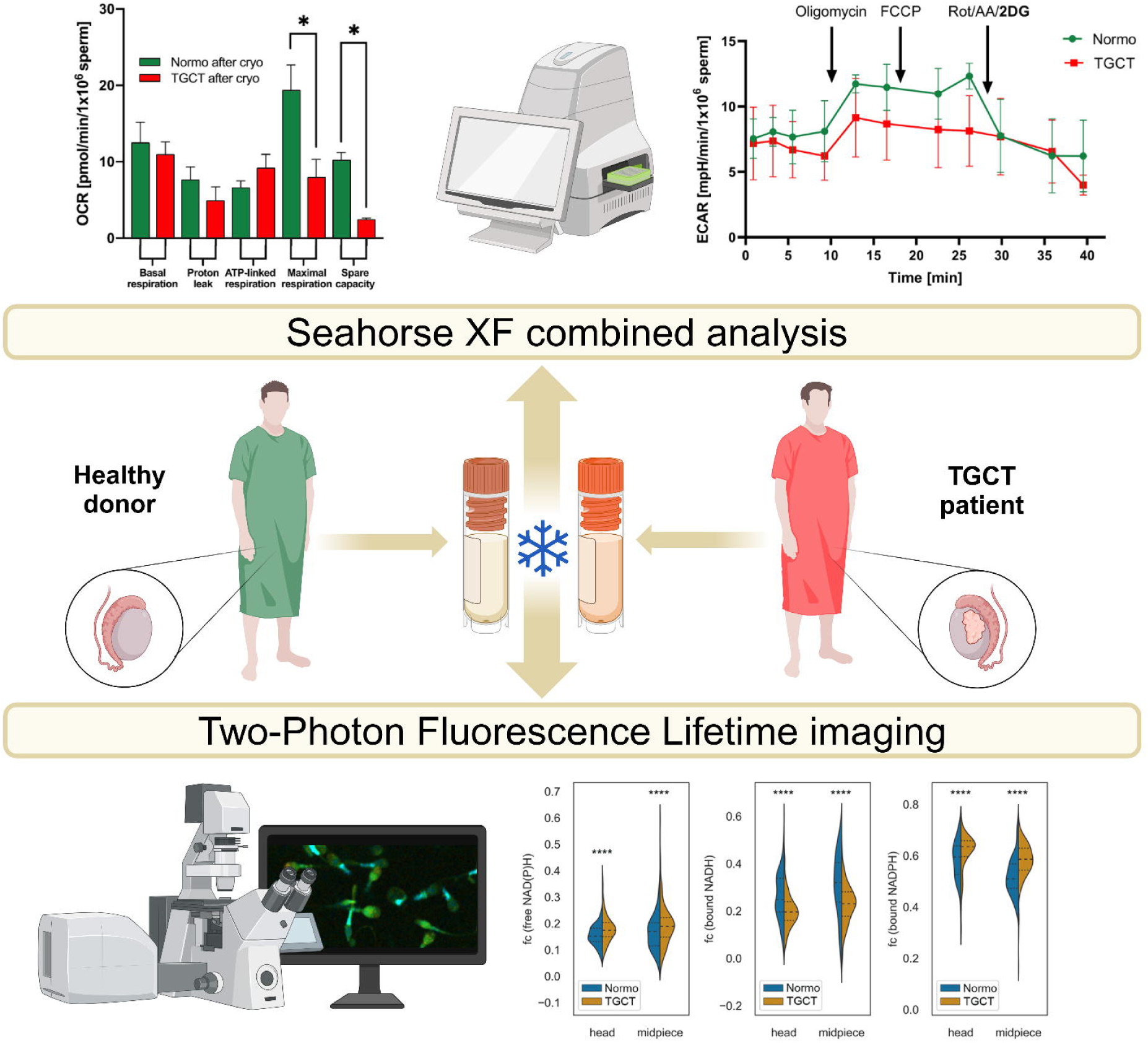

## Introduction

Testicular cancer is the most common tumour in young men between 15 and 44 years of age with steadily increasing incidence during the last few years (Krasic *et al*., 2022). Among various types of testicular tumours, testicular germ cell tumours (TGCTs), accounting for 95% of cases are prevalent (Batool *et al*., 2019, Huang *et al.*, 2022b). Since these patients undergo orchiectomy (usually unilateral), often followed by chemotherapy during their youth, maintenance of fertile sperm for future purposes by cryopreservation prior to anti-cancer therapy is the desirable approach (Djaladat *et al*., 2014). However, it has been reported that the post-thawed cryopreserved sperm samples of testicular cancer patients have the lowest recovery rate in comparison to various types of malignancies such as leukaemia, Hodgkin’s and non-Hodgkin’s lymphoma, sarcoma, brain and prostate cancers (Degl’Innocenti *et al*., 2013, Hotaling *et al*., 2013). Although the main reason has not been fully determined, mitochondrial dysfunctionality (Qasemi *et al*., 2023) may be one of the key factors contributing to low sperm vitality. In addition, there is a significant percentage of unknown, so call idiopathic, infertility and subfertility in these patient (Behboudi-Gandevani *et al*., 2021, Dias *et al*., 2020) and achieving methodological progress towards higher specificity and depth of analysis is of importance.

Spermatozoa are the most differentiated mammalian cell (Arnoult, 2020) with high-energy demand. To meet their physiological role, they require a constant efficient source of energy mainly via OXPHOS and glycolysis pathways. The degree to which these pathways are employed in sperm is species-specific and varies at different developmental and physiological stages. It has been shown that in human sperm, OXPHOS is the dominant pathway for ATP generation for spermiogenesis, maturation, and motility, while glycolysis is utilized to maintain motility and support specific functions that require quick ‘bursts’ of energy such as capacitation and the acrosomal reaction (du Plessis *et al*., 2015, Tourmente *et al*., 2022). Moreover, there is evidence that fatty acid oxidation also plays an important role in sperm physiology (Kuang *et al*., 2021). In spite of a low number of mitochondria (Hirata *et al*., 2002) and a reduced content of the cytoplasm (Irigoyen *et al*., 2022), sperm are highly metabolically active cells. This is supported by the fact that their most abundant proteins are related to energetic metabolism (Amaral *et al*., 2013).

Dysregulation of bioenergetic pathways in sperm can directly impair male fertility potential, consistent with the 17.5% of adults with infertility, which is equal to one in six couples world-wide (WHO, April 1, 2023). Out of these cases, 72% cannot be identified by current diagnostic tools and are categorized as unexplained (idiopathic) infertility (Schubert *et al*., 2019). Since mitochondria are crucial organelles in sperm not only for motility but also for protein biogenesis (Lill and Freibert, 2020), assessment of their bioenergetic efficiency can be used as a practical approach to address male infertility. In fact, this approach is important not only for fertility assessment in fresh semen but also for analysis of sperm quality after cryopreservation, which is the main strategy for fertility preservation in pathologies such as cancer and may uncover the reason behind the low recovery rate of cryopreserved sperm of TGCT patients.

In our study we used an innovative combination of tworobust methods: the Extracellular Flux Analyzer (XF Analyzer) which provides quantitative data of glycolysis and OXPHOS rates, and 2-photon Fluorescence Lifetime Imaging Microscopy (2P-FLIM) generating insights into cellular bioenergetics by analysis of reduced nicotinamide adenine dinucleotide (NADH) and its bound or unbound status. For a better insight into the principle of the method and the results it yields: NAD^+^ is reduced to NADH during glycolysis in contrary to OXPHOS where NADH is oxidized to NAD^+^. Bound NADH pool is usually relatively stable in contrary to free NADH pool (Song *et al*., 2024). Blocking or compromising the OXPHOS in mitochondria is characterized by higher incidence of shorter NADH lifetime. This can be precisely solved by multi-exponential fitting of a time-resolved fluorescence profile, an approach we used in this study (Reinhardt *et al*., 2015). As sperm after cryopreservation are very sensitive to further handling within *in vitro* conditions both methods perfectly fit to this concept as they are live-cell, real-time, minimally invasive techniques (Gómez *et al*., 2018). The XF Analyzer is routinely used in the field of cancer (Lin *et al*., 2019), immunology (van der Windt *et al*., 2016), and stem cell research (Fan *et al*., 2023). This method is important for sperm quality assessment, particularly in cases such as testicular cancer. It can deliver precise data regarding the sperm bioenergetic status in these patients. However, it should be noted that despite numerous advantages, the Analyzer has drawbacks especially due to the challenges when using pathological spermatozoa, especially after thawing. As it was shown in the study of Raad et al. (Raad *et al*., 2018), patients suffering with infertility have their functional integrity significantly affected after cryopreservation. Therefore, all sperm processing procedures have to be as careful and gentle as possible so that the quality of the spermatozoa *per se* is not secondarily affected. Since sperm manipulation and the use of different media can affect sperm viability and mitochondrial function, and as it can result in a skewed output, our initial approach was to establish a standard methodology based on the comparison between the efficiency of different media and methods using sperm from healthy normozoospermic donors before and after cryopreservation as a control group. The other technique, 2P-FLIM, was used for imaging processes associated with NADH and NADPH in live cells (Vishnyakova *et al*., 2023). This technique takes advantage of the distinct fluorescence lifetime differences between the bound and unbound forms of NADH and NADPH, offering precise details of cellular bioenergetics (Reinhardt *et al*., 2015). These coenzymes are critical to understanding cellular energy metabolism including the possibility to distinguish from which pathway the coenzymes originate (Alfonso-García *et al*., 2016). Although understanding sperm energy metabolism is pivotal for comprehending fertility and the potential treatment strategies, the use of advanced techniques such as 2P-FLIM in this domain is not as common as in other research areas. Consequently, there is lack of standardization and reproducibility across various studies, making it challenging to draw consistent conclusions from current research. Thus, we aimed to employ state-of-the-art imaging and use high-throughput methods for bioenergetic analyses of human spermatozoa originating from normozoospermic donors and patients diagnosed with TGCT before orchiectomy. With this new approach, our study delivers standardized protocols for XF analysis and 2P-FLIM as well as novel insights into the impairment of sperm motility under pathophysiological conditions caused by testicular cancer. In turn, this new approach can further help in diagnosis and personalized infertility treatment.

## Material and Methods

### Experimental design

First, due to the lack of information on the medium for evaluation of human sperm by the XF Analyzer, experiments assessing media suitability were performed. Second, the effect of cryopreservation on normozoospermic semen samples was evaluated. Third, the effect of pathology on sperm quality was assessed. For better understanding of the experimental design, its graphical representation is provided in Figure 1.

**Figure 1.**
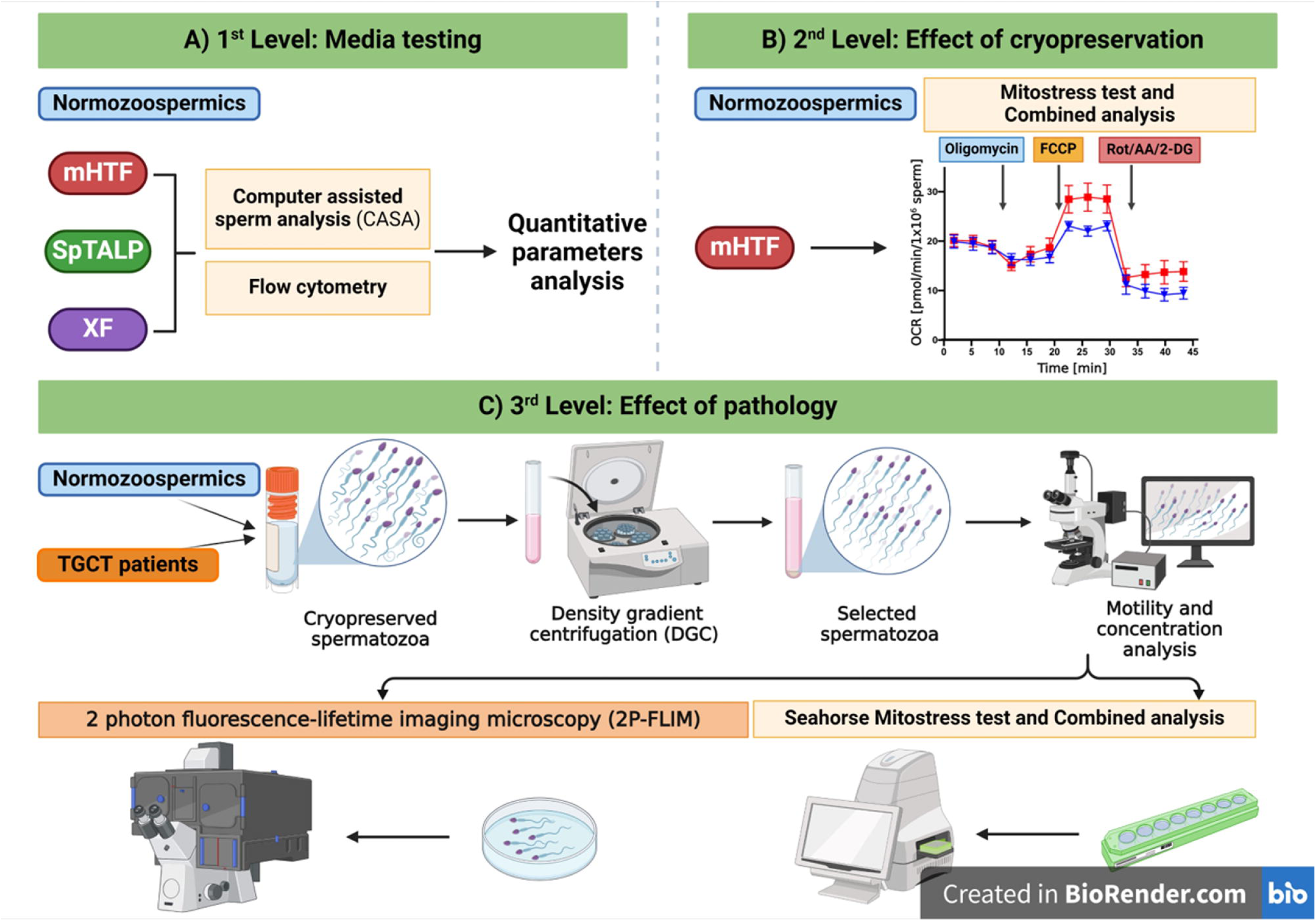
Graphical representation of the experimental design (Created with BioRender.com).

### Ethics Statement

The method of obtaining samples was approved by the Ethics Committee of the University Hospital Brno (No. 16-140421/EK), the Institute for Clinical and Experimental Medicine, Thomayer Hospital (No. A-19-09), and the General University Hospital Prague (No. 479/19 S-IV). All donors and patients signed informed consent. All samples were analysed and scored according to the WHO manual (Organization, 2010). Details from spermiograms and other patient anamnesis are listed in Supplementary Table S1. Samples from normozoospermic donors were obtained from the Centre of Assisted Reproduction of the University Hospital Brno, and those from TGCT patients were obtained from the assisted reproduction centre GENNET Ltd., Prague.

### Chemicals

All media and chemicals were purchased from Sigma-Aldrich^®^ (St. Louis, MI, USA) unless otherwise stated.

### Semen samples processing

Analysis of sperm glycolytic/OXPHOS activity was performed using samples of normozoospermic donors (n=3/n=8) and TGCT patients (n=8/n=6) before chemotherapeutic intervention. Part of the normozoospermic samples were used fresh as reference material for setting up the analysis of bioenergetic status of sperm including glycolysis/OXPHOS by the Mito Stress Test Protocol (Agilent Technologies, Santa Clara, CA, USA). Normozoospermic sperm and samples from patients diagnosed with TGCT were cryopreserved in CryoSperm (Origio, CooperSurgical^®^, Inc., Ballerup, Denmark) according to the manufacturer’s protocol. Cryopreserved samples were thawed at 37.5 °C in a circulating water bath. All sperm samples were processed by density gradient centrifugation using PureCeption 80/40 (CooperSurgical^®^, Inc.) for consequent analysis according to experimental design. In the initial part of the experimental work, the following handling and measurement media were used: modified HTF (mHTF – modified human tubal fluid) (Moody *et al*., 2017), SpTALP (Sperm Tyrode’s Albumin Lactate Pyruvate medium) and XF. Details about media composition are presented in Table 1. For insight into workflow of media preparation and whole XF analyses procedure please see Supplementary File S1.

### Computer Assisted Sperm Analysis

Analysis of sperm motility was carried out using the CASA system (ISAS, Proiser, Valencia, Spain). After pre-incubation at 37 °C for 5 min, 2 µL of the sample was evaluated in a pre-warmed 20 µm deep Leja counting chamber (in 6 different fields per sample). The recordings were made with a negative phase objective with 100× magnification and a camera with 50 FPS (frames per second) that was part of a microscope equipped with a heated stage (37 °C). On average, 200 trajectories per field were analysed. The percentage of total motility (VAP >10 µm/s) and progressive motility (VAP >10 µm/s and STR >70%) were evaluated based on the default settings of the CASA software. Results of the percentage of total motile (TMOT) and progressively motile (PMOT) sperm were obtained based on the manufacturer’s settings for human specimens.

### Flow cytometry analysis

Analyses were performed using a BD LSR Fortessa TM SORP instrument (Becton Dickinson, San Jose, CA, USA). The following lasers and filter parameters were selected, according to the used fluorescent probe: for mitochondrial integrity and viability analysis, a blue laser 488 nm (100 mW) and PE-Cy5 with excitation of 645 nm was used. The voltage was set for optimum resolution with the target of a minimum of 20,000 sperm gated events were recorded for each sample. As a standard procedure before each specific sperm parameter evaluation, positive and negative control samples were prepared to ensure the correct setting of the flow cytometer.

For viability analysis, sperm suspension (10^6^/mL) was incubated in the dark for 15 min with the novel far-red fixable viability dye DRAQ7 (ThermoFisher Scientific, Waltham, MA, USA), which stains nuclei of dead or plasma membrane-compromised sperm. Sperm mitochondrial function was assessed using JC-1 (ThermoFisher Scientific) for evaluating mitochondrial membrane potential. The final concentration of the dye was 200 nM, and the samples were incubated for 15 min in the dark. Spermatozoa with JC-1 aggregates (red fluorescence) are considered as a population with intact mitochondria with high mitochondrial membrane potential (Carrageta *et al*., 2022).

### Analysis of oxygen consumption rate and extracellular acidification rate

The Seahorse XFp Analyzer (Agilent Technologies) was used to evaluate OCR and ECAR using the standard and combined MitoStress Test assay with a modified protocol settings using the Wave software: 1 min mixing, 0.1 min waiting and 2 min of measurement. In comparison to the analysis of other cell types, inhibitor concentrations were taken from studies performed with sperm. The concentration of the uncoupler carbonyl cyanide-p-trifluoromethoxyphenyl hydrazone (FCCP) was chosen based on the titration of the optimal inhibitor level for sperm (Supplementary Figure S1). For detailed description of the whole protocol for microplate treatment, inhibitor, and media preparation see the Supplementary File S1 (Seahorse protocol).

At least 3 h before each experiment, the cartridge was hydrated with the XF calibrant solution (Agilent Technologies) in a humidified non-CO_2_ incubator at 37 °C. For each experiment, the microplate was coated with Concanavalin-A (ConA) to ensure attachment of sperm to the bottom of each well. 50 µL of the ConA working solution (0.5 mg/mL of cell culture water) was added to each well, incubated for 15 min/RT and washed two times with cell culture water.

After the remaining volume was removed, the plate was dried with air. The inhibitors oligomycin (10 µM, 20 µL), FCCP (10 µM, 22 µL), rotenone/antimycin A (Rot/AA; 5/5 µM, 25 µL) for standard MitoStress Test or rotenone/antimycin A/2-deoxyglucose (Rot; 5 µM, AA; 5 µM, 2DG; 100 mM, 25 µL) for combined MitoStress Test were pipetted into individual ports of the sensor cartridge. For negative controls (blank wells), the appropriate amount of medium used was pipetted into the corresponding ports.

After thawing and sperm selection by density gradient centrifugation accomplished using PureCeption 80/40 (CooperSurgical^®^, Inc.) according to the manufacturer’s protocol. After samples were washed 250 x g/7 min/RT in respective media and sperm motility and concentration were analysed by the CASA method. Sperm (10^6^ of sperm in 180 µL) were seeded in each microplate well and centrifuged at 200 × *g*/5 min/RT, and the plate was incubated for 10-15 min at 37 °C in CO_2_-free conditions and used for evaluation by the XFp Analyzer. Sperm ultrastructure changes in TGCT patients were imaged by Scanning Electron Microscopy according to published methodology (Poignet *et al*., 2022).

### Processing of data from XFp Analyzer

In comparison to other published studies on spermatozoa, detailed data processing techniques are provided for better reproducibility of results. The data collected by the Seahorse Analyzer were converted into the Wave software, and incorrectly evaluated wells were discarded (e.g., negative values or outliers). The Wave software allows the pO_2_ value to be checked in each well for proper oxygenation of the sample being assessed when the pO_2_ value drops after each injection and returns to the baseline value. Before exporting the data, OCR and ECAR curves for each well were checked as errors could occur by the Seahorse XFp Analyzer during the measurement process, which can be characterized by inconsistent values between wells. Inconsistent wells need to be excluded from the calculation of measured parameters. For better understanding and quality assurance of the values for OCR and ECAR derived parameter evaluation, we recommend their calculation manually in Excel or GraphPad Software according to the scheme in Figure 2.

**Figure 2.**
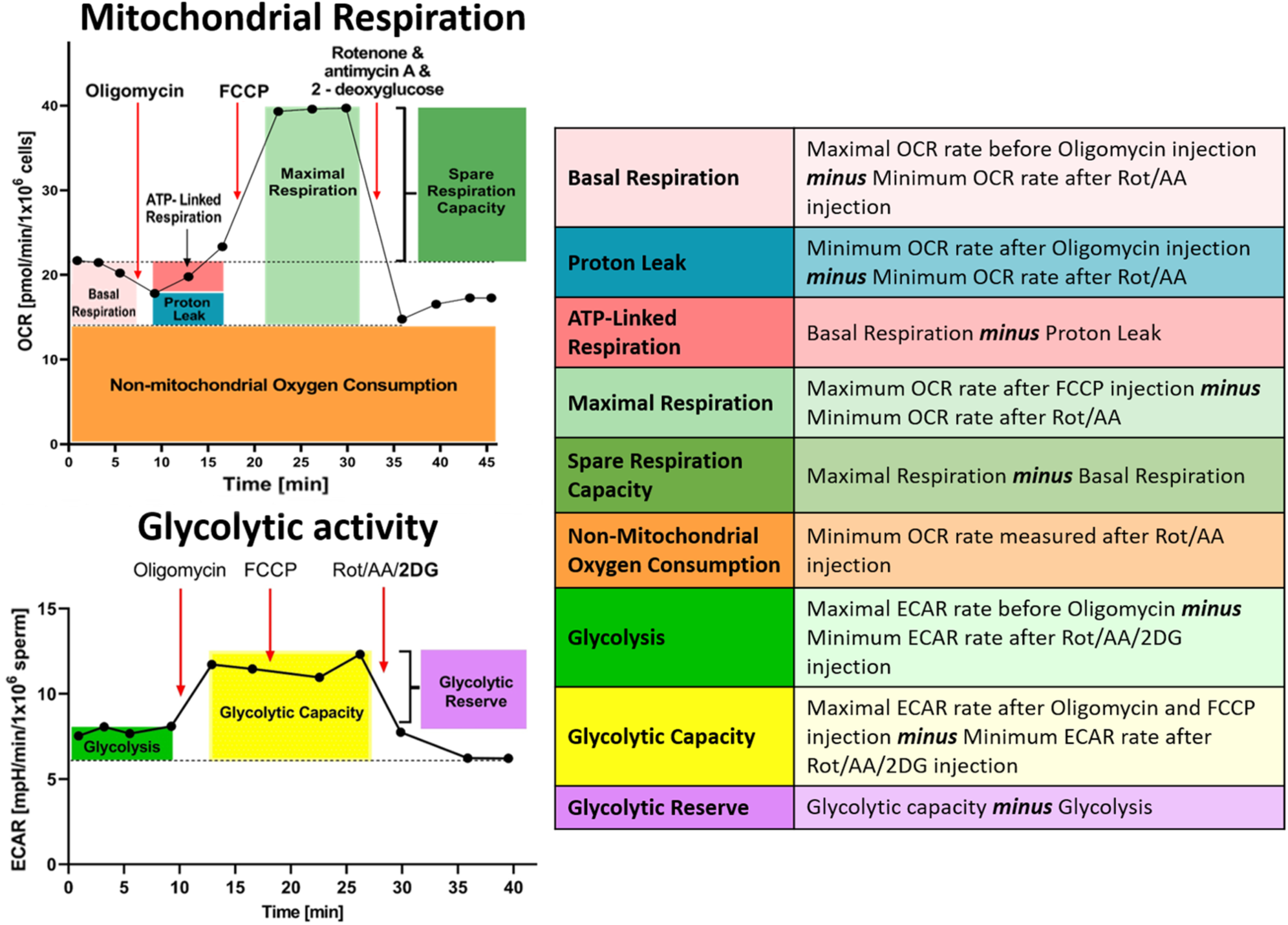
Calculation of OCR derived parameters.

Briefly, from OCR values, basal respiration was calculated by subtracting the minimum value measured after Rot/AA from the maximum value obtained before oligomycin injection. The proton leak was calculated from the minimum rate measured after oligomycin injection minus the minimum rate after Rot/AA. ATP-linked respiration is the value calculated by subtracting the proton leak from the calculated basal respiration. Maximum respiration is the equivalent value obtained from the maximum rate after FCCP injection minus the minimum rate after Rot/AA. Spare respiratory capacity is then calculated as maximal respiration minus basal respiration. Non-mitochondrial oxygen consumption is the minimum rate measured after Rot/AA injection. For calculation from ECAR values, glycolysis is the maximal ECAR rate before Oligomycin injection minus minimal rate after Rot/AA/2-DG. Glycolytic capacity is the equivalent value obtained from subtracting minimum rate after Rot/AA/2-DG from the maximal rate after oligomycin and FCCP injection. The glycolytic reserve is then calculated by subtracting glycolysis from the glycolytic capacity.

### Two-photon fluorescence lifetime imaging microscopy (2P-FLIM)

Sperm were bound to the glass bottom of a 35 mm Petri dish pre-treated with 0.7 % ConA to coat the dishes. The dish with Con-A was left for 15–20 min and then the Con-A was aspirated, and the dish was left to dry overnight under the hood. Fluorescence lifetime of NAD(P)H representing both NADH and NAD(P)H, which are mutually undistinguishable forms (Song *et al*., 2024) was assessed by 2P-FLIM using the LSM880 NLO inverted point scanning microscope (Carl Zeiss, Jena, Germany) equipped with the Chameleon Ultra II tuneable Ti-Sapphire laser (Coherent, Inc., Saxonburg, PA, USA) with 80 MHz repetition frequency. The excitation wavelength was set to 740 nm, and the sample was imaged using a 63×/1.4 Plan-Apochromat oil immersion objective with the average power of 8 mW at the sample. Fluorescence was detected by the hybrid HPM-100-40 detector (Becker and Hickl, Berlin, Germany) in non-descanned configuration. An IR blocking filter and 485 nm short pass filter were inserted before the detector to block reflected or scattered excitation light and to set the emission range to 390-485 nm. The single-photon signal counting from the detector was processed using the HydraHarp 400 Time-Correlated Single Photon Counting module (PicoQuant, Berlin, Germany) with synchronization pulses taken from the laser internal photodiode and line markers obtained from the scanner. The SymPhoTime 64 software (PicoQuant) was used for data recording and global decay fitting. A custom-made TTTR Data Analysis software was used for RGB-encoded fluorescence lifetime image reconstruction and phasor computation.

### AI machine learning

Images of sperm were segmented into individual heads and midpieces to acquire object-based excited state fluorescence lifetime features. The segmentation process was achieved in the NIS Elements software (ver._5.42.02) using a trained CNN (convolutional neural network) within the Segmentation.ai software module (available at https://doi.org/10.5281/zenodo.11277758). The network was trained on 512 × 512 pixels multichannel images containing brightfield, NAD(P)H intensity, and RGB-encoded average excited state fluorescence lifetime channels, and manually created ground truth masks were prepared. For network training, we used 59 images with the total of 1,345 midpieces, and 2,036 heads. The training was done using 700 epochs. Masks segmented using the trained network were further post-processed in the NIS-Elements and ImageJ software. Only heads larger than 80 pixels and midpieces larger than 50 pixels were included in the analysis. Additionally, intensity-based filtering of segmented objects was used. Only objects with mean NAD(P)H intensity greater than twice the background intensity of an image were included in the analysis. The background area for mean background intensity estimation was acquired by smoothing the intensity image with Gaussian kernel (sigma=3) and thresholding with Li’s method. Finally, object masks were manually checked for false positives and false negatives.

### Data analysis from 2P-FLIM

Based on the post-analysis of the acquired set of images only motile spermatozoa were subjected for further data analysis (Supplementary Video S1 as a representative image sequence). A custom-made TTTR Data Analysis software was used to process FLIM data in order to compute phasors. The NAD(P)H excited state decay histograms of each segmented object (head/midpiece) were converted to their respective phasor coordinates (G, S) with Fourier transformation. Calculation of phasor coordinates and average lifetime were done as described previously (Digman *et al*., 2008). The fit-free phasor approach yields 2D representations of the decay properties of excited states for pixel- or object-level data. This enables differentiation of groups with either similar or distinct excited state decay characteristics. When the excited state decay is mono-exponential, it falls onto the universal semicircle, while multi-exponential decays manifest within the semicircle.

### Calibration/background subtraction

Additionally, decomposition of the phasor plot using three components was performed. Overall decays from segmented images were globally fitted by re-convolution using a three-exponential model. With this, the average lifetimes of three components with mono-exponential decays representing free NAD(P)H, bound NADH and bound NADPH components were estimated. The fractional contribution of each component to the multi-exponential decays of segmented objects were extracted using the phasor approach as described previously (Torrado *et al*., 2022). Segmented objects, for which the sum of fractional contributions was larger than 1, were excluded from the analysis.

### Statistical analysis

Data of total and progressive motility (TMOT, PMOT), viability and mitochondrial integrity were analysed by one-way ANOVA, and any significant differences with the level of significance of p ≤0.05 were determined using a Tukey’s post hoc test.

Results from flow cytometry were analysed by ordinary two-way ANOVA with subsequent proof of significance using multiple comparison with Tukey’s post hoc test. Data obtained from the XF Analyzer were sorted according to the experimental design and subjected to the Mann-Whitney t-test or two-way ANOVA and Sidak’s post hoc test. All analyses were performed in GraphPad Prism (9.0.1).

Results from 2P-FLIM were analysed by non-parametric one-way ANOVA (Kruskal-Wallis test) followed by Dunn’s post hoc test to assess differences between groups.

## Results

### Selection of measuring media for sperm XF analyses

Due to the limited number of studies on human spermatozoa using the XF Analyzer and lacking experimental data regarding the various types of media for bioenergetic analysis of spermatozoa; we initially tested the most appropriate media to maintain constant level of sperm qualitative parameters. It was crucial to verify the media suitability due to the scarce sample availability. Total motility (TMOT) and progressive motility (PMOT) were not significantly affected before or after the XF analysis (Figure 3A, B). A numerical decrease in PMOT using XF media (Agilent) was evident. Based on further assessment of the sperm qualitative status in terms of viability and mitochondrial integrity, the HTF media showed the best preserving effect on sperm viability during processing/handling and evaluation. The media were finally tested during XF analysis. Results from baseline values of OCR showed constantly the highest values for the mHTF medium over the course of the protocol (Figure 3C). The effect of sperm flushing from the well and mannose presence in media on mitochondrial activity was shown not to be significant (Figure 3D). On the basis of these results, the mHTF medium was used for further sperm handling and XF analyses.

**Figure 3.**
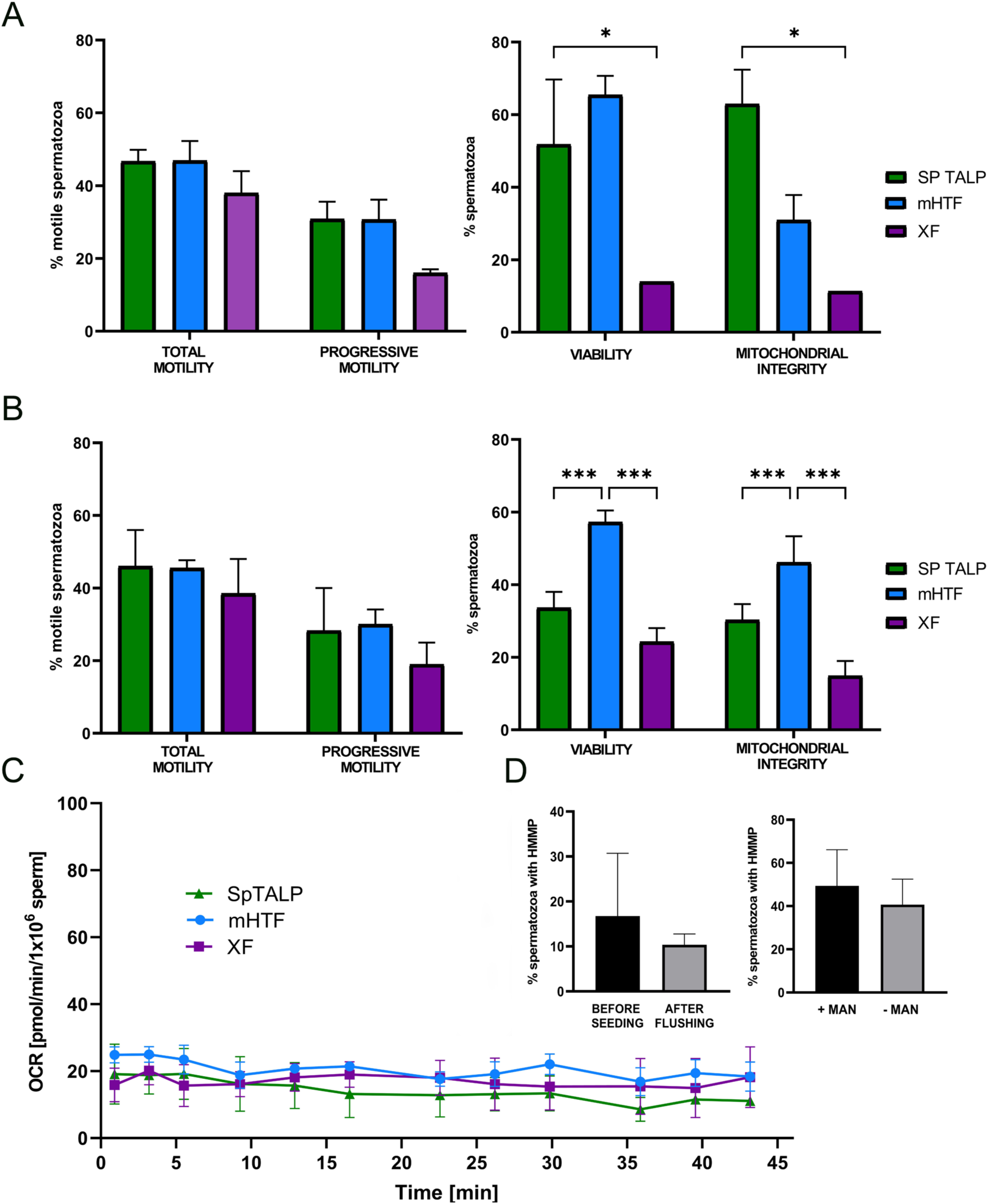
Effect of the media used for sperm handling and measuring by the XFp Analyzer on sperm qualitative parameters. A) Before and B) after analysis: total motility (TMOT), progressive motility (PMOT), viability and mitochondrial integrity; * p ≤ 0.05; *** p ≤ 0.001. C) The effect of sperm flushing from the well and mannose presence in media. D) Media effect on OCR values. Statistical tests applied were ANOVA with Tukey’s post hoc test.

### Effect of cryopreservation on OXPHOS rate of normozoospermic donors

The effect of the cryopreservation on oxidative metabolism is shown in Figure 4. Oligomycin, an inhibitor of complex V (ATP synthase) of OXPHOS, was the first injected inhibitor in the test protocol. Inhibition of complex V leads to reduced electron flow through the electron transport chain (ETC). The rate of oxygen consumption, assessed by the XFp Analyzer, decreases and ATP production by glycolysis is induced. FCCP, an OXPHOS uncoupler, was the second injected inhibitor. Its addition leads to the collapse of the proton gradient (mitochondrial membrane potential). The electron flow, which was previously inhibited by oligomycin, is restored as ETC tries to compensate the disrupted proton gradient and the recorded oxygen consumption represents maximal ETC capacity. This reading can thus give us an indication of the maximal possible respiration of a given sperm sample that could be achieved under non-physiological conditions, simulating conditions of an extreme environment. Rotenone (complex I inhibitor) and antimycin A (complex III inhibitor) were the third injected inhibitors. These agents completely suppress mitochondrial respiration and allow for subtraction of non-mitochondrial oxygen consumption, necessary for calculation of the individual parameters obtained with the MitoStress Test assay. While rates of basal respiration and ATP-linked respiration and glycolytic reserve were comparable between native and cryopreserved normozoospermic samples, the spare respiratory capacity was slightly decreased (p ≥0.05) during the process of cryopreservation/thawing influenced by the individuality of the samples (Figure 4B, C). These facts indicate that sperm bioenergetic fitness may be challenged even in healthy donors.

**Figure 4.**
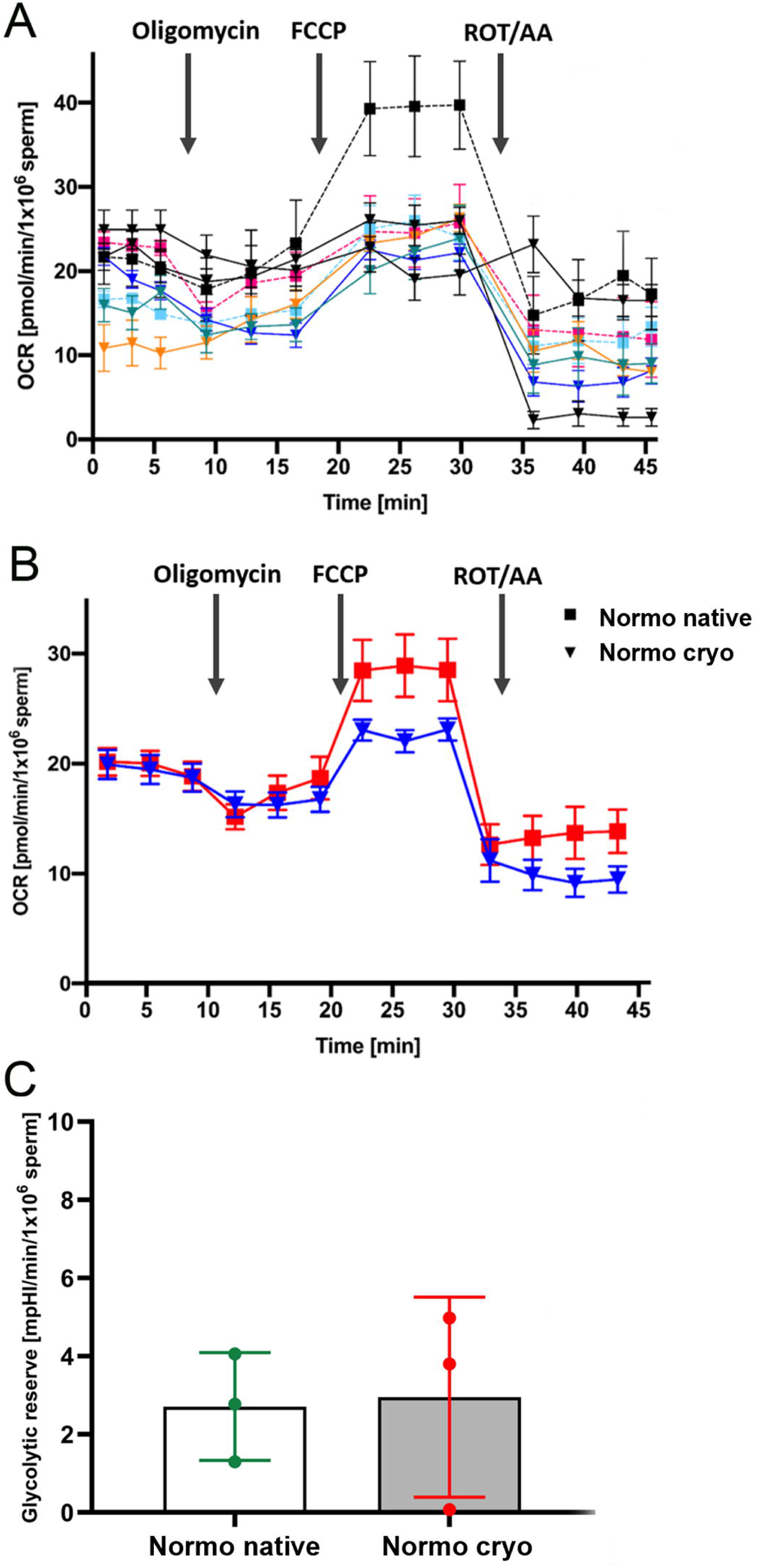
Profile of oxidative phosphorylation measured by OCR of sperm from normozoospermic donors before and after cryopreservation. A) OCR curves of individual donors; B) mean values of these two groups; C) effect of cryopreservation on glycolytic reserve parameter derived from ECAR; FCCP – Carbonyl cyanide p-(trifluoromethoxy) phenylhydrazone; ROT/AA – rotenone/antimycin A. Statistical test applied was ANOVA with Sidak’s post hoc test.

### Effect of TGCT on sperm OXPHOS and OCR derived bioenergetic indicators

Following the initial experiments using normozoospermic samples, we analysed the effect of TGCT on mitochondrial bioenergetics in six cryopreserved samples from patients with a TGCT diagnosis versus a control group of normozoospermic donors after thawing. For a better insight into the results of the analyses for normozoospermic donors and TGCT patients, the average OCR values for each group during the full course of the XF analysis are shown in Figure 5A. After cryopreservation, an overall decline in sperm bioenergetic fitness was evident in TGCT patient samples compared to normozoospermic samples. The obtained OCR in patient samples also correlates with the disruption of the sperm midpiece where mitochondria are located and OXPHOS occurs (Figure 5B). Specifically, the patient samples failed to respond to ATP synthase inhibitor oligomycin, which indicates that the impairment of sperm quality after cryopreservation is associated with the severely limited ATP-linked respiration. This negative response could also be seen in mean values (Figure 5A) (p ≥0.05) and significantly (p ≤0.05) in comparisons between individual patient samples and the normozoospermic group (Figure 5C). The response to FCCP injection was also different apart from the case of one TGCT patient compared to the control group of normozoospermic samples. The response to FCCP injection was different in one TGCT patient compared to the control group of normozoospermic samples.

**Figure 5.**
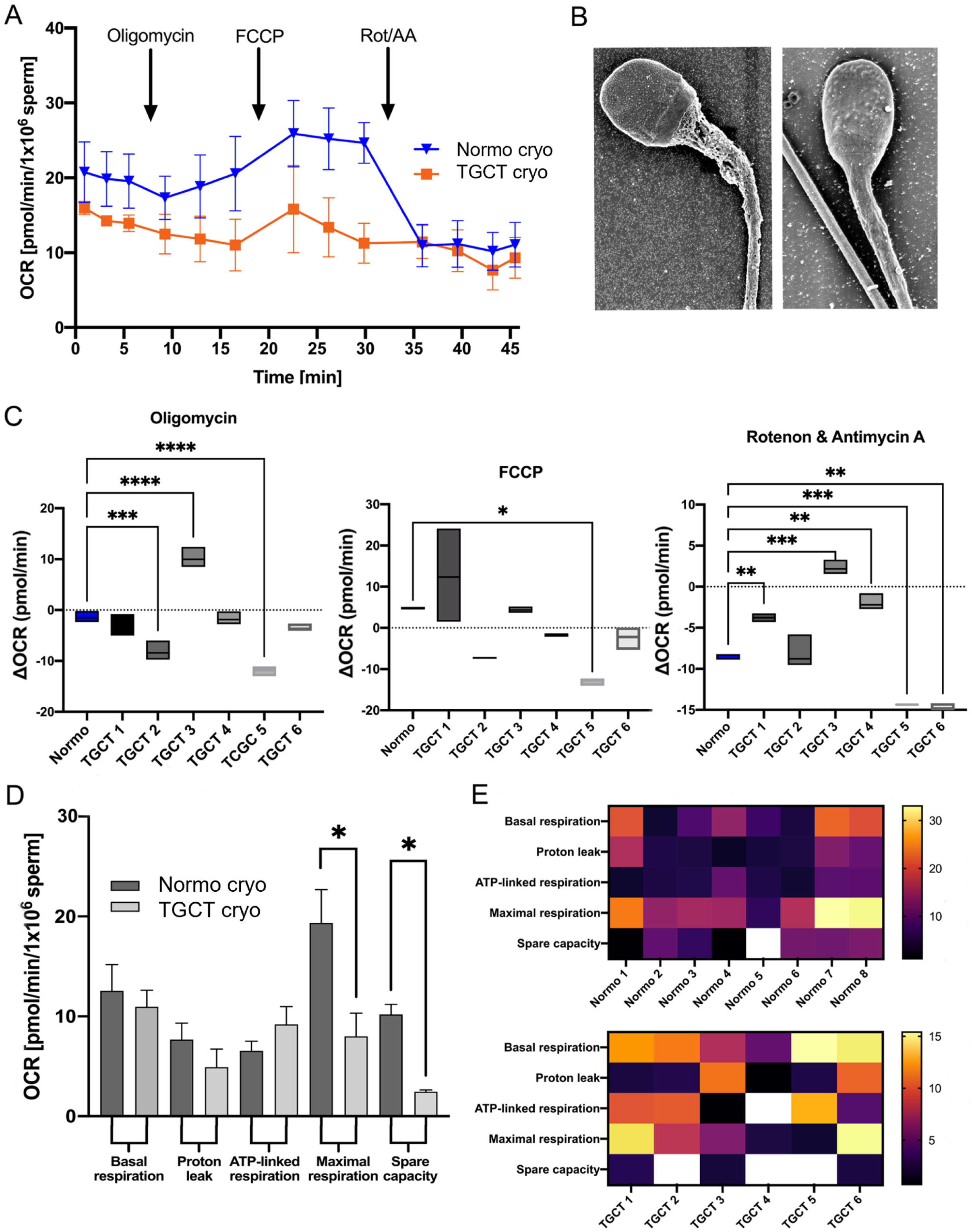
Effect of the TGCT pathology in patients before orchiectomy on the sperm oxidative phosphorylation measured by OCR during Mito Stress Test protocol. A) Profiles of OCR curves of sperm after cryopreservation from men after diagnosis for TGCT and normozoospermic donors; B) SEM representative images, on the left compromised midpiece, on the right sperm without structural abnormalities; C) Fold change (individual patients vs. mean value of normozoospermic group) of the sperm reaction on the individual inhibitors; D) OCR derived bioenergetic parameters of the sperm from normozoospermic donors and patients diagnosed with TGCT; E) Heat map representing the interindividual variability; * p ≤ 0.05; ** p ≤ 0.01; *** p ≤ 0.001; **** p ≤ 0.0001. Statistical test applied was ANOVA with Sidak’s post hoc test.

While control samples sustained the maximal ETC activity in all three recorded measurements during the course of 10 min after uncoupler addition, the TGCT sperm cells responded initially, but failed to sustain the respiratory increase compensating for unregulated proton backflow under conditions that are optimal for ETC stimulation in controls (Figure 5A). This provides further indication of compromised respiratory chain fitness in cryopreserved sperm from TGCT patients. Although individual comparison of each patient with the control group did not show a significant change in the majority of patient samples (Figure 5C), a decreasing trend (p ≥0.05) after injection of the uncoupler can be observed from the average OCR curve plot (Figure 5A). Due to the limited number of patient samples and the overall low availability of this material, titration of FCCP could not be performed for the patient group. This decreasing trend may indicate increased toxicity of this inhibitor at the chosen concentration, even though the concentration of FCCP used was titrated and considered ideal for the control group.

For increased clarity and evaluation of the data, we attempted to overcome inter-individual variability of the data by plotting them relative to the basal OCR rate value of the individual analysed samples and compare them between each other (Figure 5C). In this way, significant differences (p ≤0.0001) in the response to inhibitors in patients diagnosed with TGCT can be observed after injection of the combination of complex I and III inhibitors. This induced inconsistent decrease in OCR in the patient samples, indicating an overall reduced respiratory capacity of the spermatozoa. This may be caused by the overall reduced fitness of cryopreserved sperm due to TGCT.

As mentioned above, the MitoStress Test assay provides the possibility to calculate individual parameters that better describe the overall efficiency and the state of metabolic activity of cells. While ATP-linked respiration values were similar in normozoospermic and TGCT groups, maximal respiratory rate and spare capacity were significantly decreased (p ≤0.05) in TGCT patients’ samples (Figure 5D). These data are displayed individually for each normozoospermic control and patient sample as heat maps; for illustration of sample inter-individual variability (Figure 5E).

### Effect of TGCT on sperm glycolytic activity and ECAR-derived bioenergetic indicators

Since OCR values indicated significant differences in maximal respiration and its spare capacity (see methodology for details), we also simultaneously recorded rates of ECAR that the rate of the glycolytic pathway, of TGCT patients and normozoospermic donors. The standard MitoStress Test protocol with a specific combination of OXPHOS inhibitors allows us to obtain ECAR values representing the glycolytic reserve, a parameter indicating the ability of cells to respond to energy demand after inhibiting ATP production by OXPHOS. For the calculation of other parameters (glycolysis and glycolytic capacity) describing the glycolytic state of the cell, the MitoStress assay protocol used is insufficient (Natalia Romero, 2021). Therefore, we have introduced a combined protocol for sperm cells that allows us to correct the relevant calculation of all glycolytic parameters (glycolytic capacity, glycolytic reserve, and glycolysis) simultaneously with the measurement of OXPHOS activity by inhibiting glycolysis as the final step of the assay. Thus, by adding 2-deoxyglucose, a competitive substrate for hexokinase, we obtained background ECAR values without the actual contribution of glycolysis, which is subtracted from previously recorded rates to obtain glycolysis and glycolytic capacity parameters.

As indicated in Figure 6A, glycolysis is similarly compromised in TGCT as is mitochondrial respiration. On the other hand, there were no significant differences (p ≥0.05) in glycolytic parameters derived from ECAR values between the normozoospermic and the TGCT group (Figure 6B). Overall, Seahorse analysis of sperm bioenergetics indicated that TGCT has a more detrimental effect on mitochondrial OXPHOS than on glycolysis.

**Figure 6.**
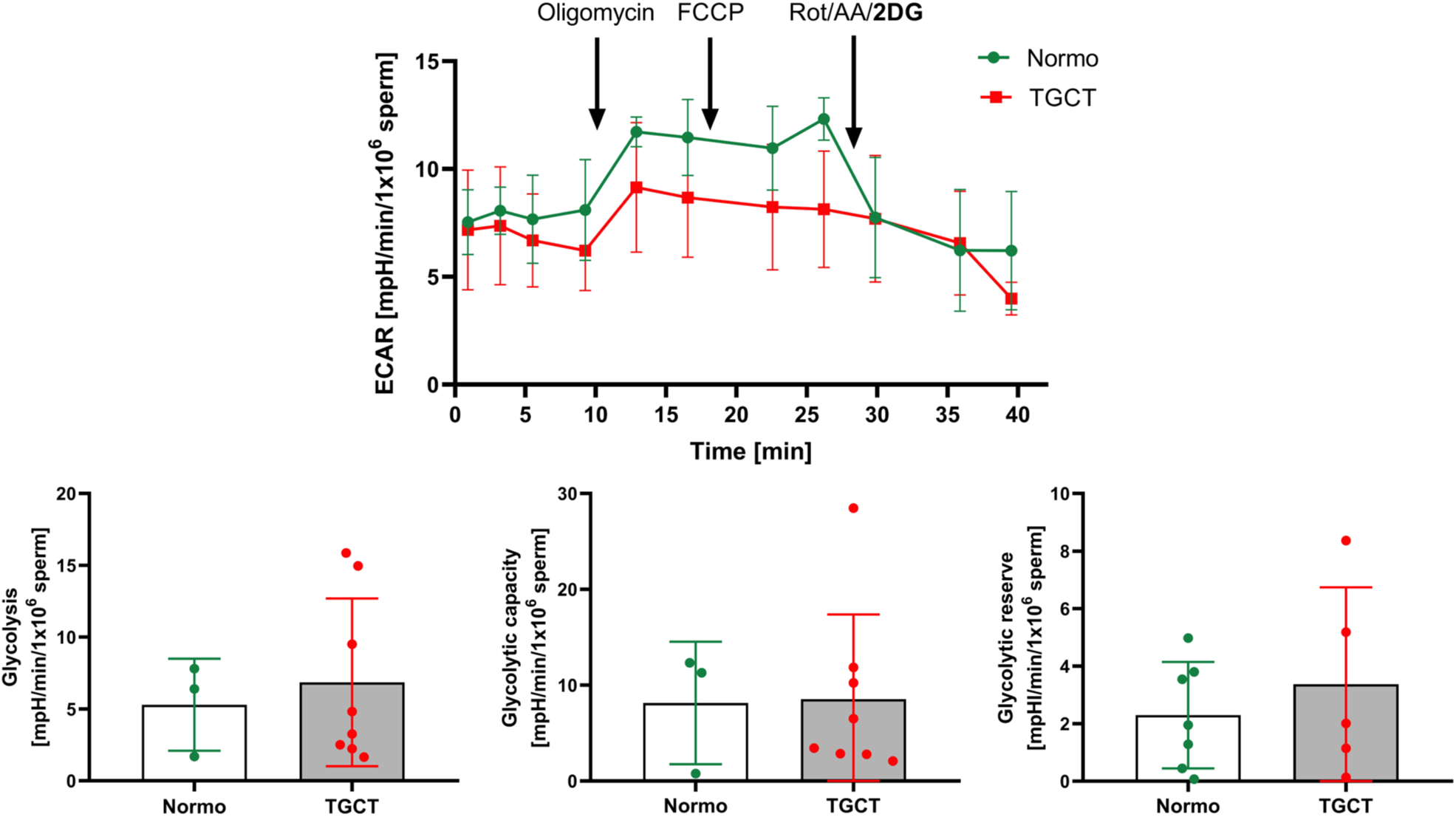
Effect of the TGCT pathology on sperm glycolytic activity in patients before orchiectomy, measured by combined Mito Stress Test protocol. A) Profiles of ECAR curves of sperm after cryopreservation from normozoospermic donors and TGCT patients; B) ECAR derived bioenergetic parameters of sperm from normozoospermic donors and TGCT patients. FCCP – Carbonyl cyanide p-(trifluoromethoxy) phenylhydrazone; ROT/AA/2DG – rotenone/antimycin A/2-deoxyglucose. Statistical test applied was ANOVA with Sidak’s post hoc test.

### NAD(P)H level in samples from normozoospermic donors and TGCT patients obtained with 2P-FLIM

All the following data are based on fluorescence images, for which detection and segmentation were performed using a customized neural network (Supplementary Figure S2). Initially, the total fluorescence intensity was analysed as a function of signal localization (sperm head and midpiece). These results are not focused on a specific form of the cofactor. Box plots in Figure 7 show distribution of the mean NAD(P)H fluorescence intensity in segmented heads and midpieces of motile spermatozoa of normozoospermic donors and TGCT patient groups. This graph shows the inter-quartile range (IQR) with the line indicating the median. Whiskers extend to points that lie within 1.5× IQR. Mean values are denoted by green triangles. In this part of the experimental work based on 2P-FLIM, significant differences (p ≤0.0001) were found in NAD(P)H signal intensity localized in sperm heads or midpieces in normozoospermic donors compares to TGCT patient samples obtained prior to orchiectomy.

**Figure 7.**
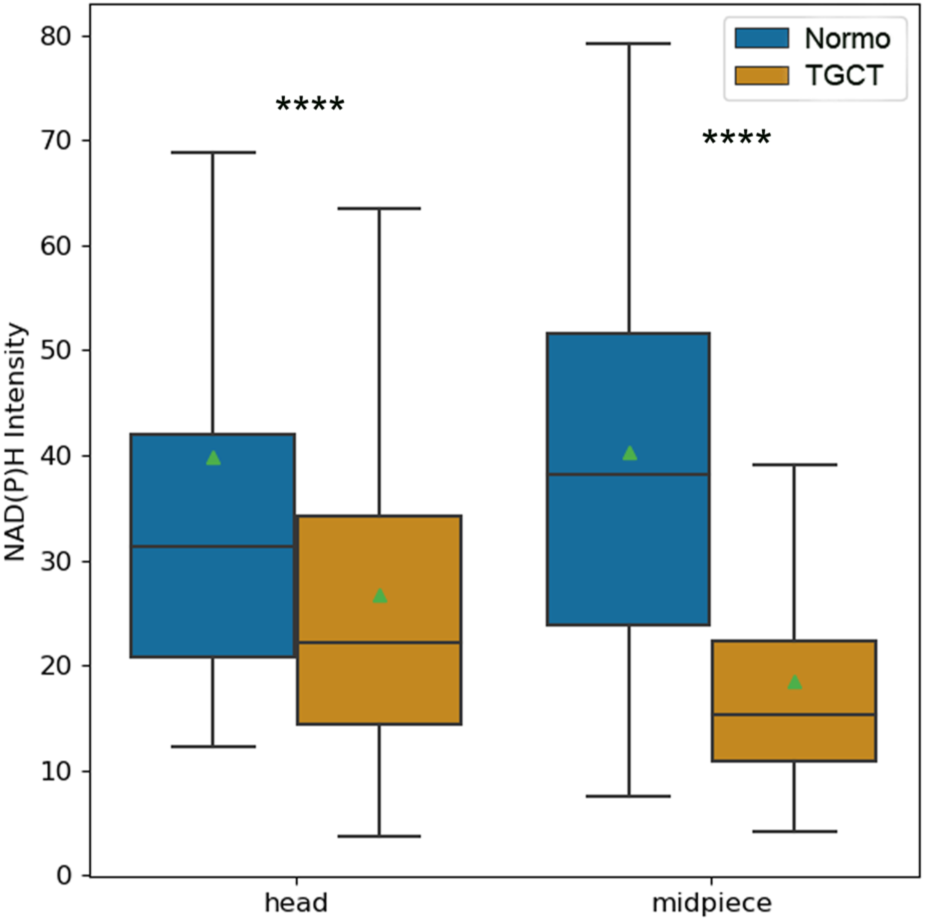
Boxplot representing signal intensity without excited state of fluorescence lifetime details obtained by two-photon fluorescence lifetime imaging microscopy (2P-FLIM) of motile spermatozoa from normozoospermic donors and patients after TGCT diagnosis and before orchiectomy; **** p ≤0.0001. The boxes depict 1st to 3rd quartile, the line ‘in the middle’ and. triangles represent median and mean, respectively. Statistical test applied was Kruskal-Wallis ANOVA followed by Dunn’s post hod test.

### Site-specific signal of NAD(P)H according to their lifetime for normozoospermic donors and TGCT patients

Phasor plots show distribution of mean fluorescence lifetimes of segmented heads and midpieces of motile spermatozoa from normozoospermic and TGCT groups acquired by 2P-FLIM (Figure 8A). Fluorescence lifetimes are plotted using the sine (S) and cosine (G) transforms to the measured decays. In the patient experimental group, the phasor cluster is shifted to the left (i.e., towards bound NAD(P)H, T = 3.53 ns) when compared to the normozoospermic group. As indicated in Figure 8B, the kernel density estimate plots illustrate the distribution of segmented heads/midpieces of normozoospermic and TGCT groups from 2P-FLIM. Contour lines mark the 10^th^, 50^th^, and 90^th^ percentiles of the data distribution. Error bars represent the standard deviation from the mean (SD). Figure 8C shows representative images of spermatozoa in both experimental groups for which the analysis was performed.

**Figure 8.**
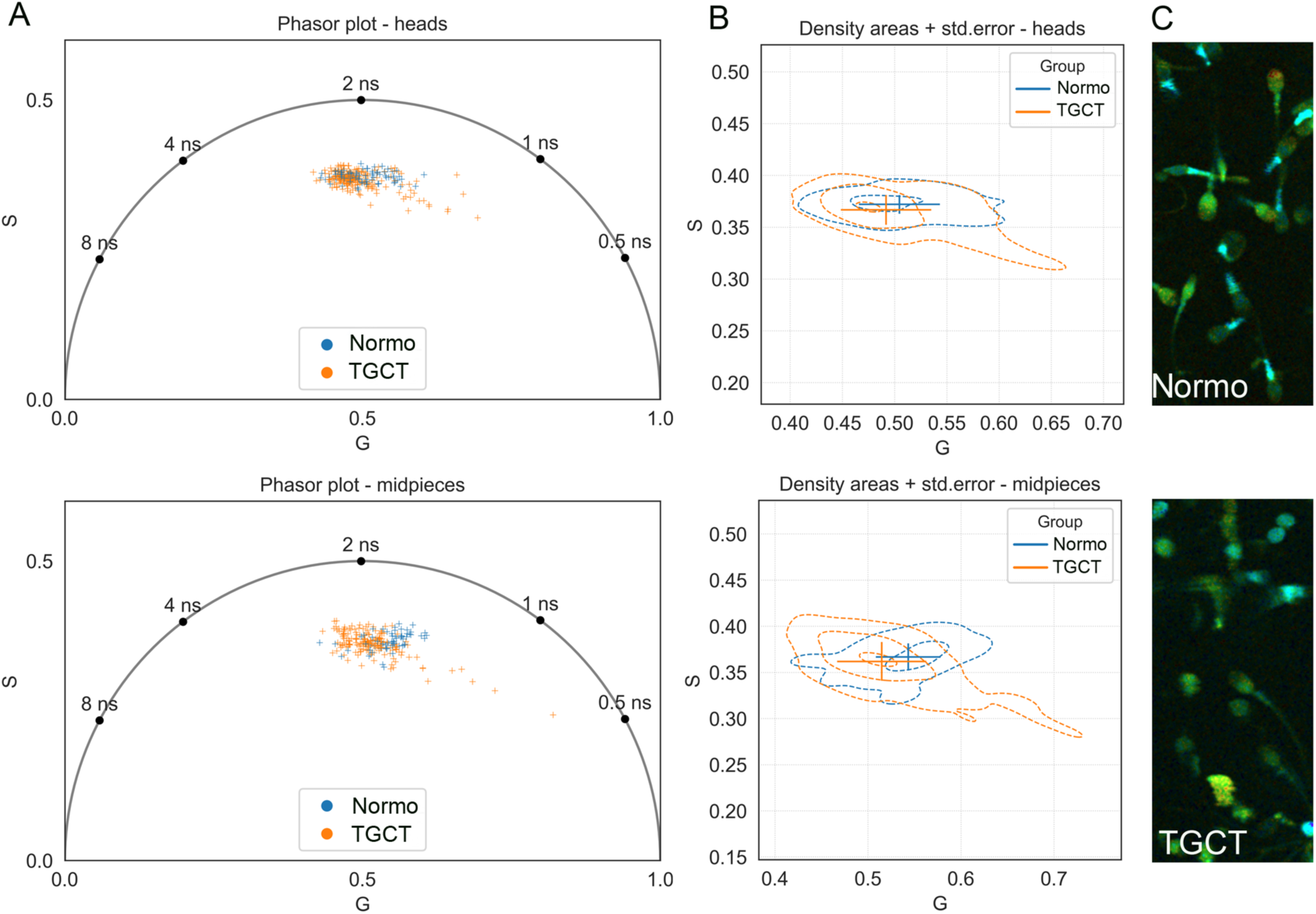
Distribution of NADH, NADPH in motile spermatozoa according to their excited state fluorescence lifetime for normozoospermic donors and TGCT patients after diagnoses and before orchiectomy. A) Phasor dot plot representation of each object (sperm/head or midpiece) related to the different lifetime components; B) Kernel density estimate plots with contour lines representing the 10th, 50th, and 90th percentiles; C) Representative images of site-specific fluorescence lifetime in sperm of normozoospermic and TGCT patients’ groups.

### Analysis of site-specific signal contribution to different lifetime thresholds

Broken-violin plots show the distribution of the mean excited state fluorescence lifetime (Figure 9A), including the fractional contributions (fc) of single-lifetime components to total intensity of segmented heads and midpieces of motile spermatozoa in normozoospermic and TGCT groups (Figure 9B). Components represent free (unbound) NAD(P)H (T = 0.27 ns), bound NADH (T = 0.95 ns), and bound NADPH (T = 3.53 ns). Dashed lines show the median, dotted lines represent the IQR range. Results of the overall lifetime in fluorescence localized in midpieces of sperm from normozoospermic donors show a significant (p ≤0.0001) shift towards lower values. Detailed analyses of the above-mentioned individual components (Figure 9B) show significant differences (p ≤0.0001) within sperm localization and between experimental groups (normozoospermic vs. TGCT patient group). Overall, NAD(P)H 2P-FLIM data indicate that TGCT mainly affects metabolic processes in the sperm midpiece, where mitochondria are localized. This finding is consistent with the Seahorse analysis, which shows downregulation of mitochondrial OXPHOS ATP production rather than changes in the glycolytic activity.

**Figure 9.**
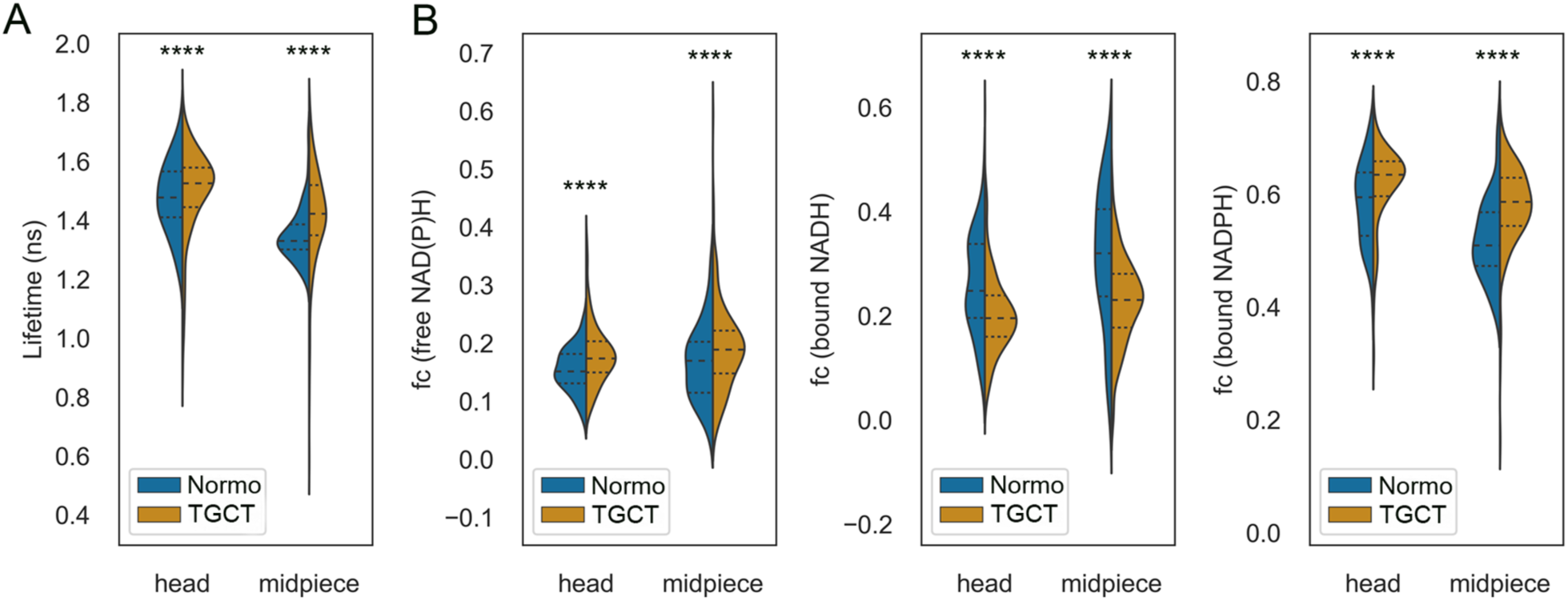
Site-specific signal of excited state fluorescence lifetime of motile spermatozoa and contribution to the different lifetime components. A) Distribution of site-specific mean fluorescence lifetime and B) fractional contribution (fc) of lifetimes from samples of normozoospermic donors and patients after diagnosis with TGCT and before orchiectomy; **** p ≤0.0001. Statistical test applied was Kruskal-Wallis ANOVA followed by Dunn’s post hod test.

## Discussion

Our research addresses the bioenergetic status associated with OXPHOS and glycolysis in cryopreserved sperm of normozoospermic donors and patients diagnosed with TGCT. As the major outcome of this study, we introduce a certain novel aspects into the protocol for precise parallel measurement of two key energetic pathways by XF Analyzer complemented by 2P-FLIM analysis. We developed an AI-based approach for detection of sperm bioenergetic status to be used for better data quantification and standardization, whereby overcoming challenges related to sperm cryopreservation and thawing (Huang *et al.*, 2022a).

Using the XF Analyzer, we present a validated methodology using HTF media with modified concentration of HEPES for sperm handling and bioenergetic analysis of human sperm samples after thawing, typically characterized by low survival rate. Our results using cryopreserved sperm samples in the HTF/HEPES modified media delivered a nearly equal level of oxidative metabolism in comparison to results of metabolic analysis of native capacitated human sperm (Freitas-Martins *et al*., 2023). These results are discussed based on data derived from sperm motility, viability, and mitochondrial integrity, and from additional experiments on the effect of media on sperm basal metabolism, when modified HTF was able to keep values within appropriate levels. The suitability of modified HTF media was consequently confirmed by the media effect on the OCR values when basal respiration reached the highest values. The bioenergetic analysis with the XFp Analyzer of cryopreserved sperm is more complicated due to the prolonged period of handling, following their exposure to different combinations of inhibitors targeting their physiological status; this finally led to readouts of the desired parameters. This was further confirmed by our results of normozoospermic samples where OCR values were affected by cryopreservation, more specifically, the respiratory reserve decreased, similar to a phenomenon observed in cryopreserved peripheral blood mononuclear cells (Korandová *et al*., 2022).

Together with the negative effect of cryopreservation, sperm quality is generally challenged by external factors such as composition of media, which represents a crucial aspect to prolonged sperm integrity, longevity, and functionality. Due to the current conditions used during the cryopreservation process, the recovery rate of human spermatozoa is strongly reduced (Freitas-Martins *et al*., 2023) and inter-individual differences are more prominent (Zhou *et al*., 2021). Therefore, establishment of our new protocols used for XF analysis compared to standard clinical practice represents a promising improvement for human sperm qualitative parameters such as viability, acrosomal, mitochondrial, and plasma membrane integrity, which are sperm features sensitive to handling procedures (Simonik *et al*., 2022). Due to specific properties of spermatozoa and the demand on the handling approach compared to other cell types, modifications of the assay medium by FCCP titration was performed to maximize the benefit of FCCP addition, which results in a proton gradient collapse (mitochondrial membrane potential dissipation). As a result, we developed a combined protocol for parallel analysis of OXPHOS and glycolysis using the XFp Analyzer. By inhibiting glycolysis as the final measurement step of this new protocol, its contribution can be specifically subtracted.

Evaluation of ECAR values provides new critical information specifically reflecting the glycolytic activity of the assessed spermatozoa. In this way, evaluation of the bioenergetic status of spermatozoa from TGCT patients and normozoospermic donors, which were used as a control group, was performed. Our results showed a significant reduction in the OXPHOS rate, particularly in reduced spare capacity and maximal respiration in sperm of TGCT patients. Despite the course of the ECAR curve being lower for TGCT patient samples, parameters derived from ECAR (glycolysis, glycolytic capacity, and glycolytic reserve) proved statistically not significantly different.

These results may indicate that TGCT is not associated with metabolic switch but rather with overall impairment of bioenergetic fitness in multiple pathways. Several studies on sperm of testicular cancer patients have revealed, using more general methodology, dysregulation of proteins involved in OXPHOS pathway for ATP production during spermiogenesis and sperm motility at the time of diagnosis (Panner Selvam *et al*., 2019). Furthermore, due to the Warburg effect, which is a phenomenon that helps tumour cells grow and which is reinforced by glycolysis, is likely dysregulated in the testis environment. Hence, alterations in bioenergetic metabolism, which we confirmed by OXPHOS and glycolysis analysis, contribute to the low sperm recovery rate after cryopreservation. Additionally, mitochondria play a role not only in ATP production but also in multiple other cellular processes (Carrageta *et al*., 2022). Fatty acid oxidation is essential for sperm fertilizing ability in mice (Kuang *et al*., 2021), and it was shown that, for example, ablation of the *Slc22a14* gene could not be compensated by another pathway to restore fertility loss.

There are metabolic studies on spermatozoa of different mammalian species (Balbach *et al*., 2023, Magdanz *et al*., 2019, Prieto *et al*., 2023), but only two studies performed metabolic profiling of human ejaculates using the XF Analyzer with subsequent correlation with basic sperm parameters (Freitas-Martins *et al*., 2023). However, so far no study has addressed the sperm bioenergetic status in testicular tumour patients, and, importantly, Freitas et al. (Freitas-Martins *et al*., 2023) analysed sperm only after capacitation, which makes it difficult to perform correlation with basic sperm parameters analysed commonly after ejaculate liquefaction. In the capacitation medium, the sperm are induced to a different metabolic status that can be dynamically changed during the capacitation period, which was not included in the published study (Freitas-Martins *et al*., 2023). Interestingly, in comparison to this study, our results of normozoospermic samples reached a higher course of OCR average values simultaneously with a similar curve. ECAR parameters obtained by the introduced protocol are of similar character to that found in their study. An additional value of our approach is the possibility to assess both energetic pathways in parallel which increases robustness of results, while reducing the input sample volume as a critical parameter in sperm samples of TGCT patients. In our study, we have also included data of correlation of calculated parameters from OXPHOS and glycolysis readouts with the basic parameters evaluated clinically in the spermiograms (Supplementary Figure S3). Parameters derived from OCR values showed the positive correlation in case of ATP linked respiration, spare respiratory capacity with total motility and progressive motility (r=0.51; r=0.48 respectively, p≥0.05). Interestingly, these results are partially consistent with the results of inferential statistics (Fig. 5D).

After identification of the changes in sperm OXPHOS and glycolysis between native and cryopreserved sperm samples and samples of TGCT patients, we investigated cofactors involved in sperm bioenergetic metabolism pathways using the 2P-FLIM method. This live-cell, real-time and label-free method for redox coenzyme imaging (Ranjit *et al*., 2019) is unique in the field of mammalian reproductive biology providing single-cell data conveying in-depth details about the cellular metabolism (Neto *et al*., 2020). In order to improve data quantification and reproducibility, we developed a novel customized assisted machine learning strategy for image segmentation and automatic detection of sperm head or midpiece. Even though motile and non-motile spermatozoa could be analysed, we included only data from motile spermatozoa in our study as relevant to previously obtained readings of OXPHOS and glycolysis. Post-processing by calibration and fitting of the acquired data with their representation in phasor plots brings further insights to the obtained results. Subsequent determination of lifetime thresholds for detailed analysis of cofactors individually allowed better understanding of data biological meaning, considering the limited knowledge regarding the redox cofactors and metabolic pathways described in the sperm (Williams and Ford, 2004). The individual components obtained by 2P-FLIM are shown within these metabolic pathways in the sperm depicted in the Scheme (Figure 10).

**Figure 10.**
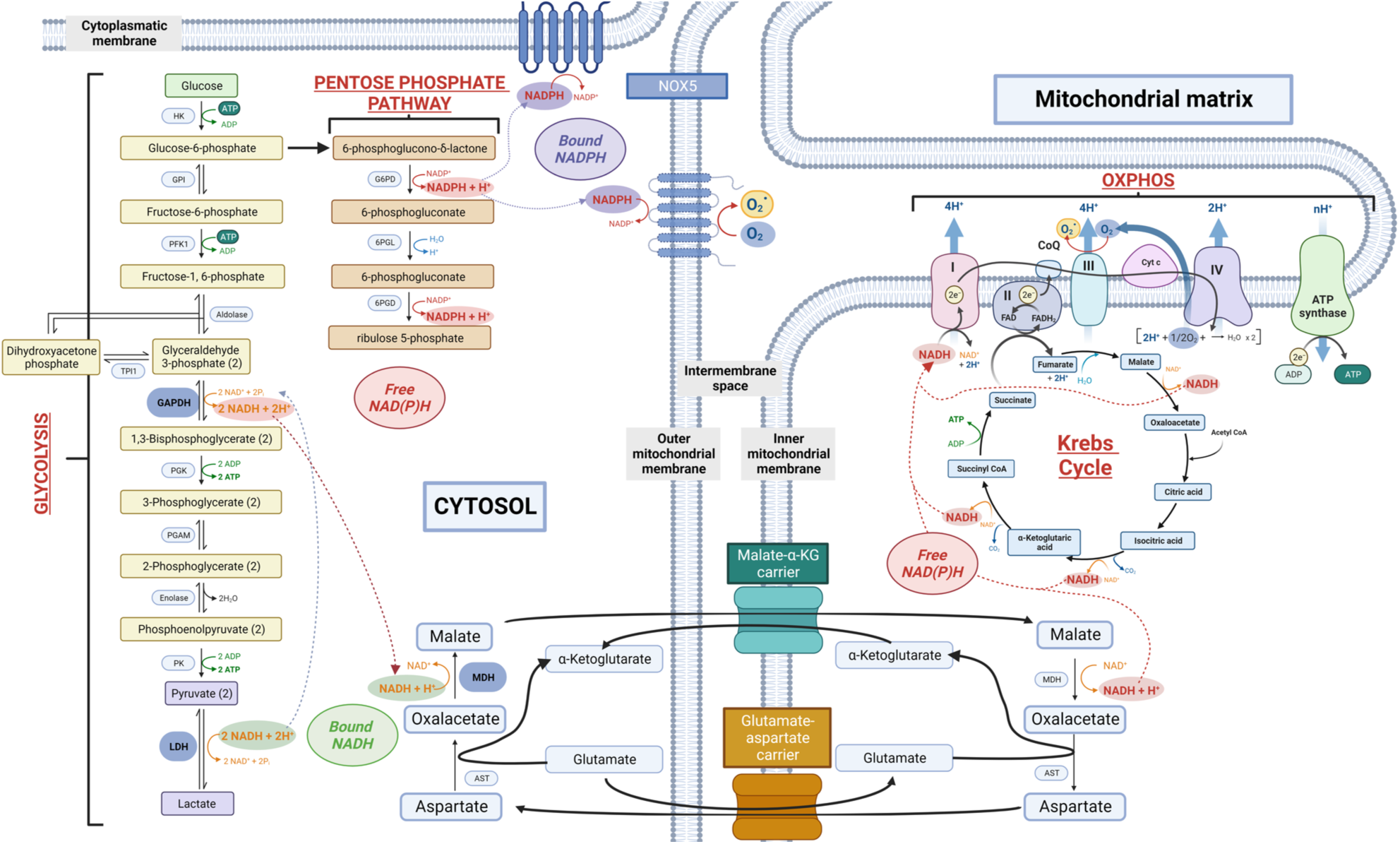
Metabolic pathways, which involve individual forms of nicotinamide adenine dinucleotide. (NADH, NAD(P)H, NADPH). Red circle shows freely diffusing NAD(P)H, green circle shows bound NADH and blue circle shows bound NADPH.

Due to the limits of this method, which cannot distinguish free NADH and NADPH from each other (Huang *et al*., 2002). The depicted scheme thus indicates combined free NAD(P)H, as it can originate in the cytoplasm from sperm glyceraldehyde 3-phosphate dehydrogenase as a cofactor for glycolysis. NAD(P)H also plays a major role in the mitochondrial matrix in the sperm midpiece as part of the Krebs cycle and is a redox cofactor delivering electrons to the ETC. The pentose phosphate pathway generates NADPH, which can also contribute to the free NAD(P)H signal. The bound form of NADH reflects NADH that is bound to lactate dehydrogenase (LDH), an enzyme of aerobic glycolysis, which regenerates NAD^+^ for further glycolysis function. Binding of NADH to LDH occurs in the cytoplasm in the course of glycolysis. However, the high activity of mammalian sperm specific LDH-C4 is essential to maintain sperm motility. The multiple localization in cytosol and mitochondria could make this enzyme a potential contributor to the replenished NAD+ formation further used in glycolysis (O’Flaherty, 2015). Another possible source of bound NADH could be NADH bound to malate dehydrogenase incorporated during the malate-aspartate shuttle. The bound form of NADPH as seen in the FLIM data, that is produced by the pentose phosphate pathway, is at higher levels in the sperm of TGCT patients. This means that it is not further processed in the cytosol and the same is true for the outer mitochondrial membrane (see the scheme in Figure 10). The differences between the experimental groups within a single localization indicate a possible higher contribution of mitochondrial damage in TGCT patients in the case of midpiece and restriction of processes within the mitochondrial membrane. Our results provide evidence that bound NADPH comprises the largest proportion of the total lifetime. This likely implies a higher relevance of this fraction for the assessment of sperm fitness. Results of the fraction in the lifetime range for bound NADH show significantly lower distribution for the TGCT patients group, which may indicate decreased metabolic activity. This is in agreement with an increase of free NAD(P)H and a shift to higher levels of bound NADPH in TGCT patients sperm heads and midpieces, which points to NADPH oxidase 5 (NOX5) activity, previously reported as an important factor in sperm cryoinjury (Keshtgar *et al*., 2020). As this enzyme influences oxidative stress balance (Miguel-Jiménez *et al*., 2021), we can hypothesize that sperm from patients are more sensitive to cryopreservation due to the effect of reactive oxygen species (ROS). In somatic cells, NOX5 localizes in membranes and in mitochondria (Touyz *et al*., 2019), and it was shown to be present in the plasma membrane in human sperm (Musset *et al*., 2012). It was shown that NOX5 activity in sperm increases due to cryopreservation and thawing. According to our results, the lifetime of the individual components cannot be fully distinguished. So far, only NADH bound to LDH is known to be present in the glycolytic pathways in the cytosol located in the sperm head and principle piece, where glycolytic enzymes have been detected (Margaryan *et al*., 2015, Miki *et al*., 2004, Yuan *et al*., 2013). Our strategy for evaluating the results based on the determination of lifetime of the components provides detailed analysis of individual cofactors, allowing better understanding of the biological context of the data, also considering the reported limited knowledge regarding the redox cofactors and pathways that occur in the sperm (Williams and Ford, 2004).

Finally, it should be mentioned that the limitation of this study lies in the relatively lower number of patient samples analysed; however, this is a factor we are unable to influence. Based on ethical principles, the limited access to patient samples was a consequence of the fact that most of them had to be used for semen banking and future use for clinical application of ART. For further follow-up studies in this or other types of pathologies, a larger number of replications need to be included to ensure that the methodological approaches presented can be further advanced into clinical practice based on more robust correlation analyses.

## Conclusion

We have set up methods required for bioenergetic analysis of human sperm based on both qualitative and quantitative approach using XF analysis and 2P-FLIM, partially controlled by artificial intelligence. These methods build on precise analysis of individual sperm samples, which represents a critical step in methodology standardization demonstrating consistent and reproducible results. We present a combined approach of XF analysis and 2P-FLIM assay to assess i) changes of oxygen consumption profile after normozoospermic sperm cryopreservation; ii) shift in patient spermatozoa in reactions to various ETC and glycolysis inhibitors; iii) site-specific variability in distribution of NADH, NAD(P)H in sperm from normozoospermic donors and from TGCT patients. For follow-up bioenergetics studies based on our results i) we recommend using a modified HTF medium for handling and measurement, and ii) we developed a trained neural network for semi-automated detection and suggest analysis of life-time decay of cofactors included in metabolic pathways.

In summary, our results show that the sperm recovery rate after cryopreservation was lowest in TGCT patients and was affected by the deterioration of bioenergetic pathways specifically in the respiratory chain and in ECAR parameters (glycolysis); this was confirmed by 2P FLIM analysis, when site-specific life-time fraction distributions were in accordance with results of XF analyses. The presented evidence points to the fact that the cryopreservation process significantly affects sperm quality, and it is enhanced in sperm of TGCT patients. We believe that the presented optimized protocol for sperm energy measurements by the XF Analyzer with an AI-based approach for 2P FLIM data processing, could be utilized for new methodological approaches in clinical practice. Implementing this sperm quality assessment via robust and precisely methodologically established methods, and identification of pathological events in cases of malignancies such as testicular cancer, could deliver benefits to the medical field and patients themselves.

## Data Availability

Source of raw data files are deposited in the link https://doi.org/10.5281/zenodo.11277540. A preprint version of this manuscript is available on bioRxiv with the DOI: https://biorxiv.org/cgi/content/short/2024.05.24.595824v1.

## Acknowledgement

We would like to acknowledge Imaging Methods Core Facility for technical assistance with the flow cytometry and scanning electron microscopy, namely Ondrej Honc and Marketa Dalecka. We thank Michala Krejci for the processing of images used for assisted machine learning and Jonathan Barlow for discussions regarding media content.

## Authors’ roles

OS experimental and graphical design, experimental work, data analysis, manuscript drafting, critical discussion; BB experimental work, data analysis, manuscript drafting, graphical design, critical discussion; VPS experimental design and work, data analysis, manuscript drafting, critical revision of final manuscript; LD data and statistical analysis, manuscript drafting, critical discussion; ZC, AB 2P-FLIM data processing, statistical analysis, critical revision of final manuscript; MQ, MF manuscript drafting; DS, ZE experimental work; TH, PS, ZK, RK, MJ, JN, LB, LZ, TB manuscript outline, critical revision of final manuscript; PPo, PPe, AP, experimental design, manuscript drafting, critical discussion; KK manuscript drafting, critical revision of final manuscript; LB, LZ, TB and KK funding acquisition.

