## Supplementary material for "Bioenergetics of human spermatozoa in patients with testicular germ cell tumour"

**List of supplementary data**

***S1 Supplementary Table:*** *Functional parameters of spermatozoa from TGCT patients (spermiograms).*

***S2 Supplementary Protocol:*** *Seahorse protocol*

***S3 Supplementary Figure 1:*** *Titration of the concentration of the mitochondrial uncoupler FCCP in relation to OCR.*

***S4 Supplementary Video:*** *Representative image sequence showing how motile sperm were detected during manual post-processing of 2P-FLIM data (*[https://zenodo.org/uploads/11235887](https://zenodo.org/uploads/11235887)).

***S5 Supplementary Figure 2:*** *2P-FLIM: Example of sperm heads and midpieces automatic detection by trained neural network.*

***S6 Supplementary Figure 3:*** *Correlation matrix representing relationship between counted parameters from XF analysis and basic semen parameters.*

***S1 Supplementary Table:*** *Functional parameters of spermatozoa from TGCT patients (spermiograms).*

| **Patient Number** | **Volume [mL]** | **Total sperm count [$10^6$]** | **Sperm concentration [$10^6$/mL]** | **Total motility [%]** | **Progressive motility A+B grade [%]** | **Morphologically abnormal [%]** | **Round cell count [$10^6$/mL]** | Sexual abstinence [days] | Liquefaction | Viscosity | Appearance | Final diagnose | Age | Medications | Smoking | Alcohol consumption | Other conditions with potential influence |
|---|---|---|---|---|---|---|---|---|---|---|---|---|---|---|---|---|---|
| **1** | 3.5 | 259 | 74 | 70.3 | 64.9 | 80 | 0 | 5 | normal | normal | normal | NORMOZOO SPERMIA | 34 | none | 18 packs/year | ocassionaly | |
| **2** | 3.5 | 157.5 | 45 | 86.7 | 80 | 85 | 0 | 4 | normal | normal | normal | NORMOZOO SPERMIA | 28 | none | no | very little | surgery for undescended right testicle in 8 years, for left varicocele in 18 years |
| **3** | 6.1 | 170.8 | 28 | 60.7 | 53.6 | 90 | 0 | 5 | normal | normal | normal | NORMOZOO SPERMIA | 29 | Alvesco | no | very little | asthma |
| **4** | 3.2 | 57.6 | 18 | 66.7 | 55.6 | 85 | 0 | 2 | normal | normal | normal | NORMOZOO SPERMIA | 35 | none | 12 packs/year | 1 beer every other day | |
| **5** | 0.7 | 98.7 | 141 | 62.4 | 61 | - | - | 3 | normal | normal | normal | NORMOZOO SPERMIA | 54 | none | occasional smoker 40 years ago | 6 times per month | |
| **6** | 4.5 | 297 | 66 | 77.3 | 74.2 | - | - | 4 | normal | normal | normal | NORMOZOO SPERMIA | 38 | none | no | very little | inguinal hernioplasty in childhood |
| **7** | 4.2 | 15.5 | 3.7 | 56.8 | 54.1 | - | - | 2 | normal | normal | gel | OLIGOZOO SPERMIA | 34 | none | no | 2 beers per day | |
| **8** | 1.5 | 303 | 202 | 31.2 | 28.2 | 92 | 5 | 10 | normal | normal | normal | ASTHENOZO OSPERMIA | 22 | none | no | no | |
| **9** | 5.3 | 106 | 20 | 60 | 50 | 90 | 0 | 6 | normal | normal | normal | NORMOZOO SPERMIA | 44 | none | no | 1 beer per day | |
| **10** | 2.5 | 122.5 | 49 | 69.4 | 59.2 | 90 | 2 | 4 | normal | normal | normal | NORMOZOO SPERMIA | 40 | none | 15 packs/year | ocassionaly | |

***S2 Supplementary Protocol:*** *Seahorse protocol.*

**MitoStress Test and Combined protocol for human spermatozoa for OXPHOS and glycolysis analyses.**

### ***I. The day before the experiment***

**Dilution of inhibitors**

First, prepare stock solutions.

| DILUTION OF INHIBITORS: STOCK SOLUTION | | | |
|---|---|---|---|
| | 8 wells plate | | |
| | stock concentration | Solvent | Mw [g/mol] |
| **Oligomycin** | 5 mM | EtOH | 802.40 |
| **FCCP** | 1 mM | DMSO | 204.62 |
| **Rotenon** | 1 mM | EtOH | 394.42 |
| **Antimycin A** | 5 mM | EtOH | 532.00 |
| **2-deoxyglucose** | 1 M | DMSO | 164.16 |

Second, prepare working solutions (see below in the section "the day of experiment").

**Media preparation**

| mHTF medium |
|---|
| **100 mL mHTF with 5 mM HEPES:** |
| **0.571 g** NaCl |
| **0.0349 g** KCl |
| **0.0024 g** $MgSO_4$ |
| **0.005 g** $KH_2PO_4$ |
| **0.0226 g** $CaCl_2$ |
| **0.1192 g** HEPES |
| **0.05 g** glucose |
| **0.0036 g** sodium pyruvate |
| **181.68 µL** sodium lactate |
| **100 µL** gentamicin |

| **100 mL Sp-TALP Stock** |
|---|
| **0.666 g** NaCl |
| **0.023 g** KCl |
| **0.0041 g** $NaH_2PO_4*H_2O$ |
| **84.9 µL** sodium lactate |
| **0.029 g** $CaCl_2*2H_2O$ |
| **0.0101 g** $MgCl_2*6H_2O$ |
| **0.119 g** HEPES (5 mM) |

| **20 mL Sp-TALP medium** |
|---|
| **20 mL** Sperm-TL stock |
| **1052 µL** sodium pyruvate (1 mM) |
| **100 µL** gentamicin (50 µg/mL) |
| **0.12 g** BSA |

| **20 mL XF medium** |
|---|
| **10 mL** 2X DMEM |
| **150.27 µL** Glucose (0.74 M) |
| **100 µL** sodium pyruvate (100 mM) |
| **195.12 µL** L-Glutamine (205 mM) |
| **200 µL** Penicilin/Streptomycin (10000 U/mL) |
| **9.354 mL** cell culture $dH_2O$ |

**Cartridge hydration**

- Ideally the evening before (12–72 h before experiment)
- Open cartridge and place the sensor cartridge with sensor plate upside down next to the utility plate
- Pipette 200 µL of XF calibrant solution to each well of utility plate, remove bubbles
- Use the XF hydrobooster (it must be removed prior to loading the inhibitors in the ports of the sensor cartridge on the day of the assay):

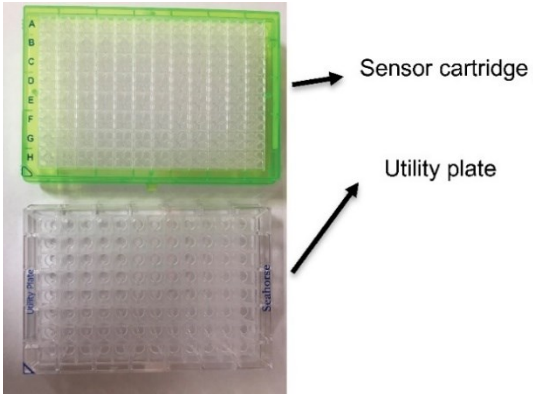


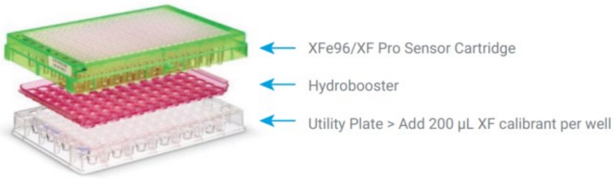


- Place in a non-$CO_2$ 37 °C incubator (if stored overnight, add a plastic beaker with water to protect the cartridge from drying).

**Cell culture coating**

**Concanavalin A** (0.5 mg/mL in $dH_2O$)

- Add 30 µL of Con A solution directly to each well and keep it for 10-15 min in RT
- Collect the rest of Con A and wash twice with cell culture water (50 µL)
- Let it dry under compressed air for 10 min (or 2 h in flowbox)

#### ***II. The day of the experiment***

- Turn on the Seahorse - MUST BE TURNED ON AT LEAST 12 H BEFORE
- Take all the media out of the fridge
  - gradient media in RT
  - assay media in 37 °C
  - remember to take coated plates out of the fridge prior seeding - must have RT
- Dilute the inhibitors

| | Concentration of working solution | Final well concentration | | |
|---|---|---|---|---|
| | | | Stock solution volume | Media volume |
| **Oligomycin** | 10 µM | **1 µM** | 3 µL | 1497 µL |
| **FCCP** | 10 µM | **1 µM** | 15 µL | 1485 µL |
| **Rotenone** | 5 µM | **0.5 µM** | 3 µL | – |
| **Antimycin A** | 5 µM | **0.5 µM** | 1 µL | |
| **2-Deoxyglucose** | 1 M | **100 mM** | 600 µL | |

**Pipetting inhibitors into the cartridge ports**

- For negative controls (blank) use the assay medium instead of inhibitors
- For cartridge calibration place the utility plate with cartridge in the XFp analyser

| | | |
|---|---|---|
| Port A | Oligomycin | 20 µL |
| Port B | FCCP | 22 µL |
| Port C | Rot/AA/2DG | 25 µL |

**Sperm sample preparation**

- Warm up the gradient media and assay media (37 °C)
- Select motile sperm cells using DGC (PureCeption, Cooper Surgical) – prepare gradient in ratio of 1:1:1 (80% : 40% : sperm)
- Centrifuge for 20 min, 300 x g, brake 0, RT
- Remove first two layers of gradient and then take only the pellet
- Check the sperm concentration and motility in the pellet
- You need 1 million sperm per well
- Calculate the amount of sample for dilution (1 million of sperm in 180 µL of media for 1 well)

**Sperm cells seeding and running the seahorse protocol**

- Dilute the original sperm suspension with the assay medium (pre-warmed to 37 °C) to the final concentration of 1 million sperm in 180 µL of the suspension (= 1 well)
- Use wells in the corner for BLANK – assay medium without sperm
- Centrifuge the plate for 5 min at 200 x g at RT, check motility using inverted microscope
- After seeding put the plate to 37 °C for 10 min (use non-$CO_2$ 37°C incubator)
- When calibration of the utility plate with cartridge is completed, remove the utility plate from the machine and place the cell plate on the tray

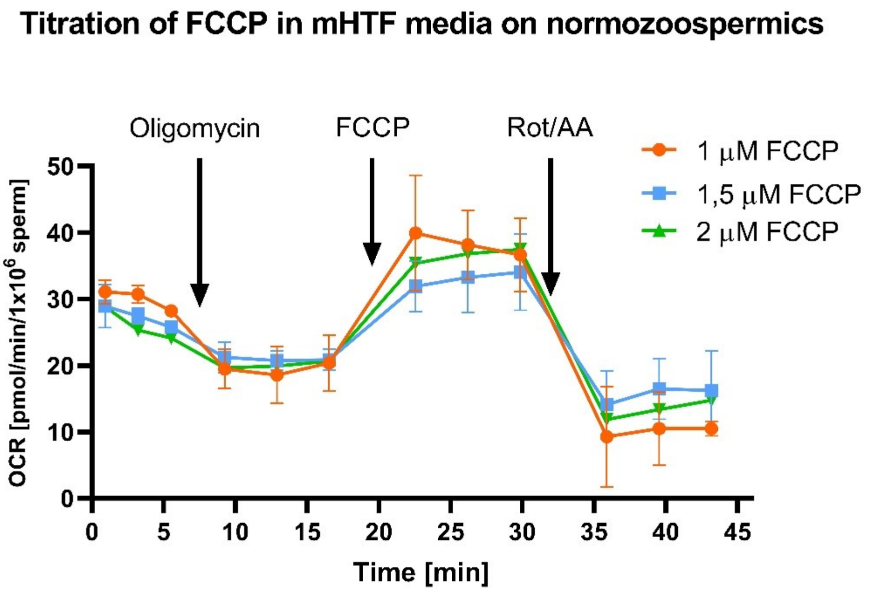


***S3 Supplementary Figure 1****: Titration of the concentration of the mitochondrial uncoupler FCCP in relation to OCR.*

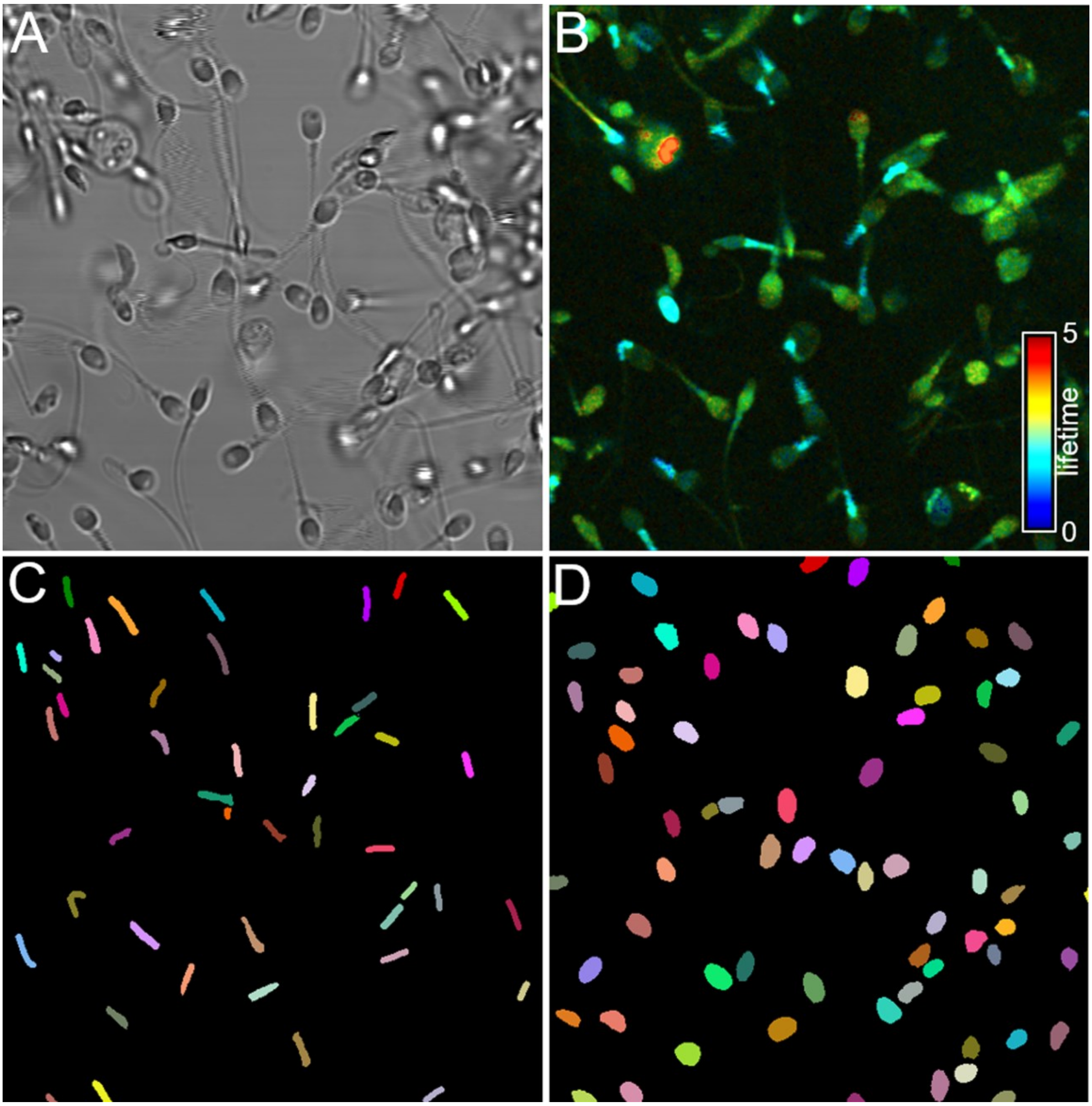


***S5 Supplementary Figure 2:*** *2P-FLIM: Example of sperm heads and midpieces automatic detection by trained neural network.*

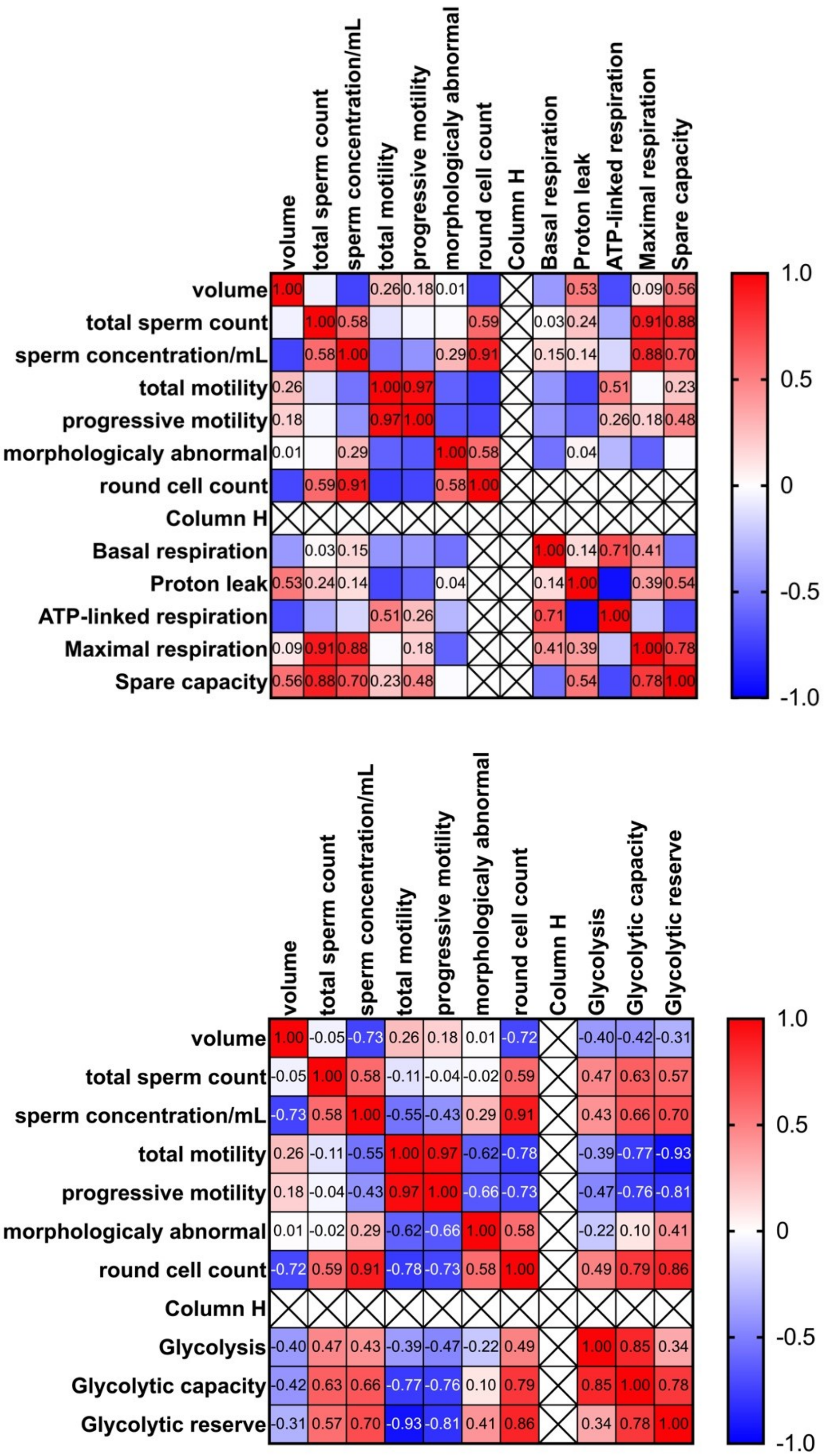


***S6 Supplementary Figure 3:*** *Correlation matrix representing relationship between counted parameters from XF analysis and basic semen parameters.*
